# Acute particulate matter (PM_10_) exposure selectively triggers behavioral alterations in the presymptomatic Experimental Autoimmune Encephalomyelitis (EAE) mouse model of Multiple Sclerosis

**DOI:** 10.1101/2025.03.20.644303

**Authors:** Martino Bonato, Francesca Montarolo, Roberta Parolisi, Maura Caudana, Giuliana Abbadessa, Vita Cardinale, Giulia Nato, Claudio Pandino, Niccolò Di Cintio, Valentina Schiavo, Maurizio Giustetto, Federico Luzzati, Silvia De Francia, Antonio Bertolotto, Annalisa Buffo, Enrica Boda

**Author notes:** **<u>Corresponding author:</u>** Enrica Boda, Department of Neuroscience Rita Levi Montalcini, Neuroscience Institute Cavalieri Ottolenghi (NICO), University of Turin, Regione Gonzole, 10 – 10043 Orbassano (Turin), Italy. equal contribution (Martino Bonato, Francesca Montarolo).

## Abstract

Multiple Sclerosis (MS) is a chronic disease of the Central Nervous System, where neuroinflammation and autoimmune response against myelin lead to functional impairments, cognitive and psychiatric symptoms. Exposure to air pollution – in particular to peaks of particulate matter (PM) – has been associated with an increase of hospital admissions for MS onset and relapses and exacerbated neuroinflammation in MS patients. Here, in the MOG_35-55_-induced experimental autoimmune encephalomyelitis (EAE) mouse model of MS, we tested the hypothesis that exposure to PM_10_ might influence the disease course and severity in individuals with a predisposing background. Short-term PM_10_ exposures - occurring either before immunization or during the pre-symptomatic phase - did not modify disease manifestation in EAE mice, as assessed by clinical and neuropathological analyses. Yet, presymptomatic EAE – but not healthy - mice selectively showed increased disinhibited, risk-taking and novelty-seeking behaviors early after being exposed to PM_10_. These data show a selective vulnerability of immunologically primed mice toward the effects of PM_10_, occurring before the emergence of overt motor impairment and presenting as specific behavioral alterations.

## Introduction

The pathogenesis of autoimmune diseases, including Multiple Sclerosis (MS), is thought to have a strong genetic background component, but a substantial impact of environmental risk factors – including exposure to air pollution, tobacco smoking, infections or drugs – has been also recognized in either the development or the exacerbation of these conditions (Sellner et al., 2011; Gawda et al., 2017). MS is a chronic autoimmune disease of the Central Nervous System (CNS) - characterized by neuroinflammation, multifocal demyelination, and eventually neurodegeneration - leading to sensory alterations, motor and autonomic impairments and, in most cases, psychiatric and cognitive symptoms (e.g. Woo et al., 2024; Benedict et al., 2020; Sparaco et al., 2021). These latter disturbances present as depression or anxiety, reduced memory, slower information processing and executive functions. Deficits of attention and alterations of decision-making abilities have been also reported, with an important impact on the quality of life and daily activities in people with MS (Paul et al., 1998; Toro et al., 2018; Neuhaus et al., 2018).

Studies have linked exposure to air pollution, and, in particular, short-term increases in airborne particulate matter (PM), with an increased incidence of hospital admissions for MS onset and relapses (Angelici et al., 2016; Roux et al., 2017, Heydarpour et al., 2014; Oikonen et al., 2003; Gregory et al., 2008; Noorimotlagh et al., 2021). This suggests that even acute exposures to high levels of PM might accelerate the onset of MS in individuals with a predisposing background or trigger the worsening of the disease in MS patients (Bergamaschi et al., 2021).

Despite recent improvements in air quality, air pollution is still the largest environmental health risk in Europe, where more than 80% of the urban population is exposed to unsafe concentrations of PM, according to the World Health Organization (WHO) Air Quality Guidelines (Europe’s air quality status 2024 released by the European Environment Agency ; https://www.eea.europa.eu//publications/europes-air-quality-status-2024). PM is a mixture of particles - typically including inorganic compounds, aromatic hydrocarbons (e.g. benzene), metals, and microbial components (e.g. lipopolysaccharide; Becker et al., 2002) - released from the combustion of fossil fuels, gasoline, diesel, coal, or wood, as well as from natural events like volcanic eruptions, soil and rocks erosion (Sierra-Vargas et al., 2012). PM particles are categorized by their size, with coarse particles (i.e. PM_10_) and fine particles (i.e. PM_2.5_) having aerodynamic diameters smaller than 10 and 2.5 μm, respectively. Thanks to their size, PM particles can penetrate the respiratory tract up to the alveoli. Here, PM initiates a local oxidative-inflammatory reaction that eventually extends and disrupts the homeostasis of various distant organs and systems, including the CNS (Peeples, 2020). Consistently, even in healthy subjects, residency in cities with high air pollution including PM, is associated with neuroinflammation, white matter injury, disruption of the blood brain barrier (BBB) integrity and altered brain innate immune activity (Calderón-Garcidueñas et al., 2008; Babadjouni et al., 2017). Such a response appears to be amplified in MS patients, where PM exposure correlates with brain inflammatory exacerbation as detected by MRI with gadolinium (Bergamaschi et al., 2017). Also, of relevance for MS, higher concentration of antibodies directed against myelin proteins have been found in the serum and in the cerebrospinal fluid (CSF) of people resident in metropolitan city areas with high levels of airborne PM (Calderón-Garcidueñas et al., 2015). Besides this correlative evidence in humans, controlled studies in experimental animal models have revealed that PM exposure causes neurovascular inflammation, BBB functional deficits (Nejad et al., 2014), microglia activation and demyelination (Woodward et al., 2017; Han et al., 2022). We also found that acute exposure to PM targets the endogenous regenerative capability of the CNS tissue by hampering remyelination and promoting astroglia and microglia reactivity in the mouse (Parolisi et al., 2021). More recently, by investigating the micro-RNA (miRNA) cargo of plasmatic extracellular vesicles (EVs) in an animal model of MS (i.e. the chronic MOG_35-55_-induced experimental autoimmune encephalomyelitis - EAE), we also showed that an acute PM_10_ exposure triggers a selective deregulation of miRNAs targeting transcripts coding for MS-relevant factors involved in nervous system-, immune- and inflammation-related pathways, possibly contributing to disease worsening (Bonato et al., 2025).

Thus, there is now a strong rationale to hypothesize that even short-term exposure to PM can interact or even synergize with the individual’s immune background and contribute to MS pathogenesis or exacerbation. To address this issue, we investigated whether an acute exposure to PM_10_ - occurring either before immunization or during the pre-symptomatic phase - affects the disease course and the behavioral phenotype of the MOG_35-55_-induced EAE mouse model of MS. While acute PM_10_ exposures did not significantly accelerate the emergence of motor deficits, nor modified the disease course in EAE mice, presymptomatic EAE – but not healthy - mice selectively showed increased disinhibited, risk-taking and novelty-seeking behaviors after being exposed to PM_10_. To assess the biological substrate of the selective response of EAE mice to PM_10_, we investigated possible alterations of the dopaminergic neurotransmission, which were formerly associated with a behavioral phenotype reminiscent of that observed in PM_10_-exposed EAE mice (Pogorelov et al., 2005; Kalueff et al., 2016). Yet, no change in dopamine availability or in the expression of genes coding for dopamine receptors, transporters and synthetic/degrading enzymes have been detected as associated with PM_10_ exposure in the whole brain as well as in dissected brain regions relevant for the observed behaviors (i.e. striatum and prefrontal cortex). Overall, data show a selective vulnerability of immunologically primed mice toward the effects of PM_10_, occurring before the emergence of overt motor impairment and presenting as specific behavioral alterations.

## Materials and Methods

### Animals and experimental design

Mice were housed in the vivarium under standard conditions (12-hr light/12-hr dark cycle at 21°C) with food and water ad libitum. The project was designed according to the guidelines of the NIH, the European Communities Council (2010/63/EU) and the Italian Law for Care and Use of Experimental Animals (DL26/2014). It was also approved by the Italian Ministry of Health (authorization 510/ 2020-PR to EB) and the Bioethical Committee of the University of Turin. The study was conducted according to the ARRIVE guidelines.

### Chronic EAE induction and evaluation of the clinical score

To induce chronic EAE, 8 week-old female C57BL/6J mice (Charles River, Calco, Italy) were immunized by 2 subcutaneous injections of 200 μg myelin oligodendrocyte glycoprotein 35-55 peptide (MOG35–55; Espikem, Florence, Italy) in incomplete Freund’s adjuvant (IFA; Sigma-Aldrich, Milan, Italy) containing 8 mg/ml Mycobacterium tuberculosis (strain H37Ra; Difco Laboratories Inc., Franklin Lakes, NJ, USA), followed by 2 intravenous injections of 500 ng of Pertussis toxin (Duotech, Milan, Italy) on the immunization day and 48 h later (Montarolo et al., 2015). Body weight and clinical score of each individual mouse were recorded daily by an operator blind to the mouse treatment (i.e. PM *vs.* saline, see below) for 21 days. The clinical score was designed to evaluate the differences in the onset and course of the pathology following the severity of motor defect symptoms, and the score was set as follows: 0=healthy; 1=limp tail; 2=ataxia and/or paresis of hindlimbs; 3=paralysis of hindlimbs and/or paresis of forelimbs; 4=tetraplegia; 5=moribund or dead. Median clinical scores were calculated for each group per day to analyze the disease course of the EAE. Cumulative and maximum score, as well as the percentage of disease-free (score=0) mice were also calculated.

### PM_10_ administration

We exposed healthy (Ctrl) and EAE mice to a commercially available PM_10_ (NIST Standard Reference Material 1648a; Sigma-Aldrich), which is a chemically standardized compound widely used in toxicological studies (Akhtar et al., 2013; Kewcharoenwong et al., 2023; Gałuszka-Bulaga et al., 2023). Stock suspensions of NIST SRM 1648 PM were prepared in sterile ultrapure water and aliquots were thoroughly mixed under sonication for 1 h prior to each experiment. Mice were randomly divided into control (saline) and treatment (PM) groups. The treatment group was treated by intratracheal instillations (Intubation stand, Kent Scientific Corporation) of a PM_10_ suspension (10 μg in 50 μl of saline), and the control group was treated with saline (50 μl), as in our previous study (Parolisi et al., 2021). We opted for the intratracheal administration of PM to assure low variability of absorption and to exclude a direct nose-to-CNS transit of PM via the olfactory mucosa.

PM_10_ dose was calculated considering the daily respiratory volume of mice (0.04 m^3^) and the daily peaks of PM_10_ concentration in East Europe polluted areas (>250 μg/m^3^; Zibert et al., 2016). Therefore, in order to use a PM_10_ dose relevant for human exposure, the dose used in this study was set as 10 μg (0.04 m^3^/day × 250 μg/m^3^). Two different protocols of acute exposure were adopted: a pre-immunization protocol, where PM_10_ was administered 3 days and 1 day before EAE immunization; and a post-immunization protocol, where PM_10_ was administered 4 and 6 days after EAE immunization. Both protocols assured administration of PM_10_ during the presymptomatic phase of the EAE model, where debilitating symptoms are yet to be displayed by the animals.

### Behavioral Tests

Tests were performed 6 days after EAE immunization (i.e.in the presymptomatic phase), 6 hours after the second exposure to PM_10_ or saline to both EAE and Ctrl mice. Animals were habituated to the testing room in single transparent cages (36 × 21 × 14 cm height) for at least 1 hour before the test. Analysis was performed by an operator blind to the treatment of the animals, and mouse performance was video recorded with the operator outside the testing room.

### Open field (OF) test

Locomotor activity was investigated by means of the OF test, under dim white light conditions (2 lux). Each animal was placed in the corner of the arena (50 × 50 cm) and let explore the arena for 1 h under videorecording. Total distance and distance traveled in the center (25 × 25 cm) of the arena were analyzed using Ethovision XT video track system (Noldus Information Technology, Wageningen, The Netherlands, RRID:SCR_000441).

### Analysis of the stereotypic behaviors

Video recordings obtained during the OF test were also analyzed to highlight patterns of stereotypic behaviors, such as grooming (i.e. cleaning of the fur of the head/snout with forepaws, adjusting of posture) and rearing (i.e. extension of the mouse forelimbs when standing on the hindlimbs). Using the open-source Behavioral Observation Research Interactive Software (BORIS, www.boris.unito.it, RRID:SCR_025700) for behavioral quantification, each stereotypic behavior was measured for total events (i.e. number of time the animal performed the action) and total time (i.e. seconds or minutes spent in total by the animal performing a single behavior).

### Elevated Plus Maze (EPM) test

The EPM test was used to evaluate anxious behaviors. The apparatus used for the test is a plus-cross shaped platform made in gray forex and raised 60 cm above floor level. The platform comprises two open arms (30 × 5 × 0.20 cm) and two closed arms (30 × 5 × 15 cm walls) originating from a central square platform (5 × 5 cm). At the beginning of each trial, each mouse was gently placed on the center of the square platform and allowed to explore the maze for 5 minutes under videorecording. The number of entries and the cumulative time spent in either open or closed arms was measured with the Ethovision XT software (RRID:SCR_000441). Animals were considered to have entered an arm of the maze when all four paws left the center of the square.

### Novel Object Location (NOL) Test

The test comprised three distinct phases: a first phase of habituation with an empty enclosed arena (33 × 33 cm); a second phase, where two identical objects have been placed into the arena (each object was situated near the corners of one side of the arena); and a third phase, where one of the two objects has been moved to the opposite side of the arena. Inter-trial interval was 10 minutes. During each phase, the animal could freely explore the arena for 10 minutes under videorecording. Mouse movement and interaction with the objects have been tracked using the Ethovision XT software (RRID:SCR_000441).

### Tail suspension test

Mice were suspended by the tail into an apparatus (20 × 40 × 60 cm) with white walls and an open side to be video recorded. The animals were attached to the apparatus by adhesive tape wrapped at 1 cm from the tip of the tail. A solid white divisor allowed for simultaneous testing of two mice. Each trial lasted for 6 minutes, and immobility time of each mouse was manually measured (immobility was determined as absence of struggles/movement for >1s). Percentage of time of immobility and latency to first immobility were used to assess depressive symptoms.

### Histological Analyses

At 21 days post-immunization (dpi), animals were deeply anesthetized (ketamine-xylazine 90 mg/kg and 9 mg/kg, respectively) and transcardially perfused with 4% paraformaldehyde (PFA) in 0.1 M phosphate buffer (PB). Brains and spinal cords were postfixed overnight, cryoprotected, cut in 30 μm thick sections, and processed according to standard immunohistological procedures (Boda et al., 2022). Sections of the lumbar enlargement of the spinal cord were collected in PBS and then stained to detect the expression of different antigens: MOG (myelin oligodendrocyte glycoprotein; 1:1000, Proteintech Cat# 12690-1-AP, RRID:AB_2145527) to visualize myelination; IBA1 (ionized calcium-binding adapter molecule 1; 1:1000, FUJIFILM Wako Pure Chemical Corporation Cat# 019-19741, RRID:AB_839504), as a microglia marker; GFAP (glial fibrillary acid protein; 1:1000, Agilent Cat# GA524, RRID:AB_2811722) as a marker of astrocytes. Incubation with primary antibodies was made overnight at 4 °C in PBS with 1% Triton-X 100. The sections were then exposed for 2 h at room temperature (RT) to secondary Cy3 (1:500, Jackson ImmunoResearch Labs Cat# 711-165-152, RRID:AB_2307443) or Alexa Fluor 488 (1:500, Jackson ImmunoResearch Labs Cat# 711-545-152, RRID:AB_2313584)/Alexa Fluor 647 (1:500, Jackson ImmunoResearch Labs Cat# 711-605-152, RRID:AB_2492288) -conjugated antibodies. After processing, sections were mounted on microscope slides with Tris-glycerol supplemented with 10% Mowiol (Calbiochem, LaJolla, CA).

### Image processing and data analysis

Histological specimens were examined with a Zeiss Axioscan Z.1 Slide Scanner, which allowed precise tile-scanning of whole-section images. Images were acquired at 20× magnification (Plan-Apochromat 20×/0.8) with a pixel size of 0.325 × 0.325 μm. Quantitative evaluations were performed on images with ImageJ (Research Service Branch, National Institutes of Health, Bethesda, MD; available at http://rsb.info.nih.gov/ij/; RRID:SCR_003070). To analyze the expression level of MOG, IBA1 and GFAP in the lesion area, the positive fractioned area (i.e. the percentage of positive pixels throughout the entire lesioned area) was quantified. Six animals/group and 10-15 sections per animal were analyzed for each experimental condition. Adobe Photoshop 6.0 (Adobe Systems, San Jose, CA, RRID:SCR_014199) was used to assemble the final plates.

### Spectral confocal reflectance imaging and analysis

Spectral confocal reflectance microscopy (SCoRe), which takes advantage of the high refractive index of myelinated tissue (Schain et al., 2014), was used to detect and quantify compact myelin using reflected light, as described in (Craig et al., 2024). Reflectance images of myelin in 4,6-diamidino-2-phenylindole (DAPI, to counterstain cell nuclei) -stained histological specimens were captured on a Leica Stellaris confocal microscope (Leica Microsystems) with an acoustic-optical beam splitter (AOBS) with a 70/30 reflectance/transmission (RT) ratio. To obtain reflection images we selected 499 and 633 wavelengths with AOBS set in reflection mode and we selected narrow detection bandwidths centered around the 499 and 633 nm lasers, in order to obtain excitation light reflected by the specimen as the detection light. Confocal images (1024 × 1024 pixels) were acquired as z-stacks (voxel sixe x,y = 0.658 μm; z=1μm) at 20×, with a speed of 200 Hz and 56.6 μm pinhole size. For quantitative analysis, the reflection signals from both laser lines were merged into a single composite image.

Compact myelin signal was assessed by positive pixel identification using a minimum threshold cutoff with the Otsu plugin in ImageJ (Research Service Branch, National Institutes of Health, Bethesda, MD; available at http://rsb.info.nih.gov/ij/; RRID:SCR_003070). To generate a percentage, the area positive for SCoRe signal was divided by the total area of the region of interest (ROI). In all cases, the ROI was a trace made to include the total area of the grey matter and white matter in each section analyzed. 4-5 animals/group and 3-7 sections per animal were analyzed for each experimental condition. Adobe Photoshop 6.0 (Adobe Systems, San Jose, CA, RRID:SCR_014199) was used to assemble the final plates.

### Quantification of dopamine by High Performance Liquid Chromatography (HPLC) and High Performance Liquid Chromatography-High Resolution Mass Spectroscopy (HPLC-HRMS)

Dopamine (DA) concentration was measured in the whole brain tissue as well as in the prefrontal cortex (PFC) and striatum of Ctrl and EAE mice sacrificed 6 hours after exposure to saline or PM_10_. Dissected samples were stored at −80°C until use. DA hydrochloride and DA-d4 hydrochloride (analytical standards), formic acid and ammonium formate (both HPLC grade) were purchased from Sigma-Aldrich (Milan, Italy). Methanol (MeOH) and acetonitrile (ACN) (both HPLC grade) were purchased from VWR (Milan, Italy). HPLC-grade water was produced by a Milli-DI system coupled with a Synergy 185 system by Millipore (Milan, Italy).

For DA quantification in the whole brain, HPLC analysis was performed as reported in (Montarolo et al., 2022). Specifically, previously weighted brain samples were immersed in liquid nitrogen, sonicated for 1 min, reconstituted in 1 ml of water, and sonicated for another min. The calibration curve of DA was established in the concentration range of 0.01–5.00 μg/ml. 100 μl brain samples were extracted by protein precipitation using 300 μl of freeze solution of formic acid 0.5%v/v in acetonitrile. Each sample was mixed for at least 15″ and then stored in freezer at −20°C for 15′ and later centrifuged at 4000 rpm for 10 min. 250 μl of supernatant were transferred to an injection vial. Chromatographic separation was performed at 35°C, using a column oven, on a RP column (Atlantis T3 4.6 × 50 mm, 5 μm, Waters, USA. The flow rate was set at 1 ml/min.

For DA quantification in dissected brain areas, to enhance method sensitivity, HPLC-HRMS methodology was used. Samples were sonicated in 500 μl of water and subsequently mixed with 500 μl of H_2_O/MeOH (1:1) with 0.025 M formic acid, vortexed (10 s), and centrifuged (3000 rpm, 10 min). Supernatants were loaded on pre-conditioned SPE columns, washed twice with H_2_O/MeOH (2:8) with 0.025 M formic acid, and eluted with H_2_O/MeOH (2:8) containing 0.050 M formic acid and 0.060 M ammonium formate. Eluates were dried at 45°C in a UniVapo centrifuge for 2h, resuspended in 200 μl of 0,2% formic acid in water and transferred to microvials for HPLC-MS. DA levels were quantified by HPLC-HRMS using a timsTOF Pro 2 (Bruker Daltonics, Bremen, Germany) equipped with a Luna Polar C18 column (2.1 × 150 mm, 3 μm; Phenomenex, Bologna, Italy) in binary gradient mode with solvent A (0.2% formic acid in H_2_O) and solvent B (0.1% formic acid in MeOH). The gradient was: 0 min, 5% B; 8–9 min, 100% B, 9,1-12 min 5% B. Flow rate was 0.2 ml/min. The HRMS was operated in positive ion modes with VIP-HESI source (end plate offset 500 V, capillary voltage 4.5 kV, nebulizer 2 bar, dry gas 10 L/min, dry temp 230 °C, sheath temp 400 °C). Full-scan acquisition (50–200 m/z) was performed at 30,000 resolution (FTMS). Auto MS/MS was used for DDA, with collision energy of 20 eV (50–200 m/z), acquiring spectra at 16–20 Hz in profile mode. Data were processed with TASQ-2025b (Bruker).

In both cases, DA concentrations were expressed as μg/mg tissue. The method’s limit of detection (LOD) and limit of quantification (LOQ) were optimized using DA and DA-d4 standard solutions.

### Quantitative RT-PCR

RNA extraction, reverse transcription and quantitative Real Time RT-PCR was performed as described in (Boda et al., 2022), either with predeveloped Taqman assays (Applied Biosystems, Thermofisher, Waltham, USA) or by combining the RealTime Ready Universal Probe Library (UPL, Roche Diagnostics, Monza, Italy) with the primers listed in Supplementary Table 2. Real Time data were collected on the Applied Biosystems StepOnePlus Real-Time PCR System with StepOne™ Software (RRID:SCR_014281). Data analysis was performed with Microsoft Excel (Microsoft Office 365, RRID:SCR_016137). A relative quantification approach was used, according to the 2-ddCT method. β-actin was used to normalize expression levels.

### Statistical Analyses

Statistical analyses were carried out with GraphPad Prism 9 (GraphPad software, Inc, RRID:SCR_002798). The Shapiro-Wilk test was first applied to test for a normal distribution of the data. When normally distributed, unpaired Student’s t-test (to compare two groups) and Two-ways ANOVA test (for multiple group comparisons) followed by Bonferroni’s post-hoc analysis, were used. Alternatively, when data were not normally distributed, Mann–Whitney U-test (to compare two groups) was used. The Kaplan-Meier test was used to compare percentages of disease-free mice over time. In all instances, P *<* 0.05 was considered as statistically significant. Graphs represent individual data and mean ± standard error of the mean (SE). Statistical differences were indicated with * P *<* 0.05, **P *<* 0.01, ***P *<* 0.001, ****P *<* 0.0001. The list of the applied tests and number of animals in each case are included in Suppl. Table 1.

## Results

### Acute exposure to PM_10_ does not modify EAE disease course

To validate PM exposure as a contributing factor for disease onset/worsening in an animal model of MS, we have combined the induction of chronic EAE in mice and PM_10_ exposure. Specifically, to assess whether PM operates either by facilitating the induction of autoimmunity (thereby contributing to the initiation of the pathology) or by exacerbating an already established pathological setting (thereby operating as a secondary stressor), mice were acutely exposed to PM_10_ (or saline, as a control) either before the immunization (Fig.1**a**) or during the pre-symptomatic phase (Fig.2**a**), respectively. Daily assessment of the mouse clinical score and body weight, performed for 21 days after EAE immunization, showed that acute PM_10_ exposures did not accelerate the emergence of motor deficits, nor modified the disease course in EAE mice (Fig. 1**b**,**c**; Fig. 2**b**,**c**). The percentage of disease-free mice over time as well as maximum and cumulative EAE scores were also not significantly affected by pre- and post-immunization PM_10_ exposures (Fig. 1**d**-**f**; Fig. 2**d**-**f**).

**Figure 1.**
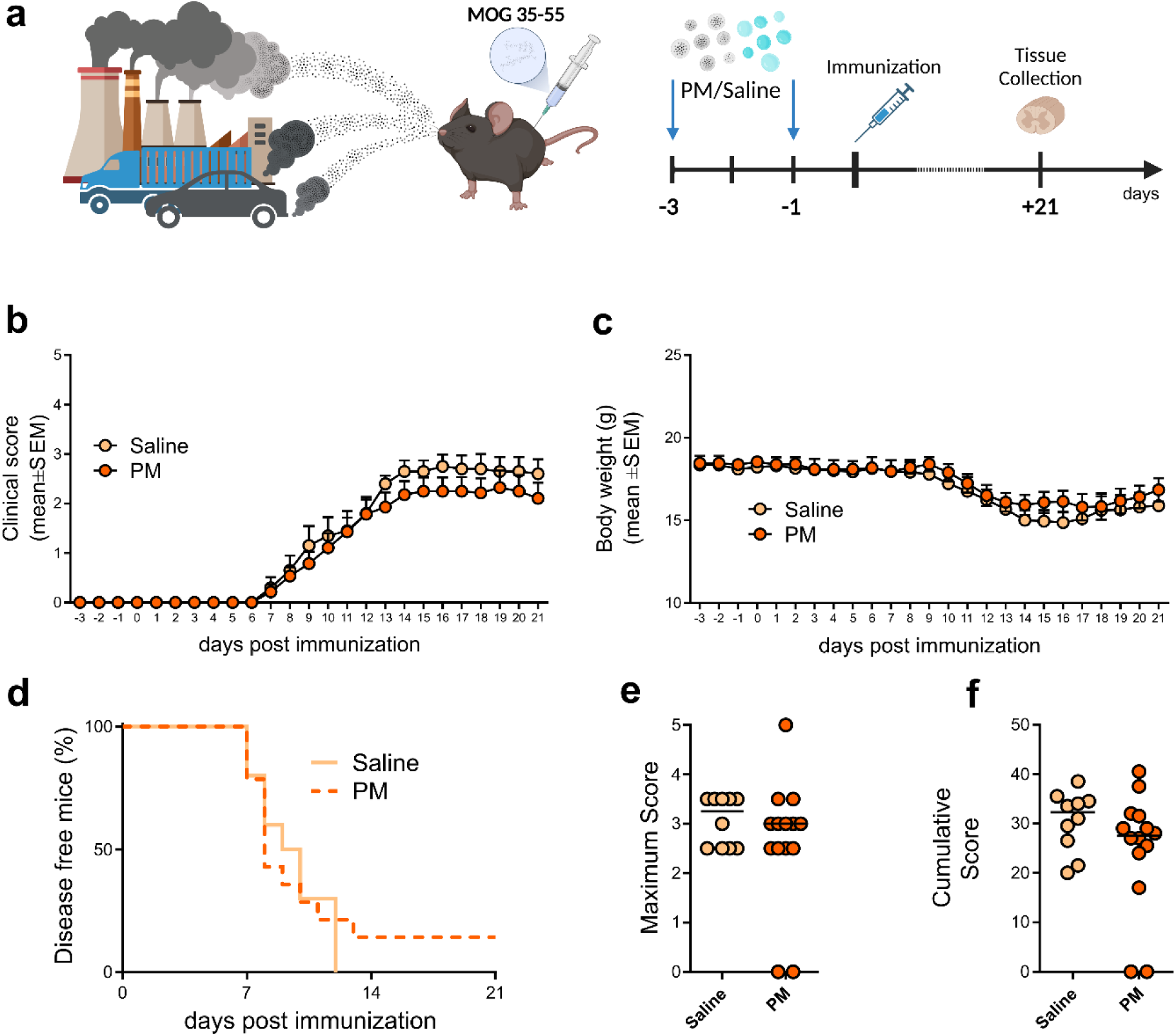
Pre-immunization exposure to PM_10_ does not alter EAE pathological course. (**a**) Schematic representation of the experimental design (graphics created with BioRender.com). (**b**) Mean clinical score of saline vs. PM_10_-exposed EAE mice. (**c**) Mean body weight of saline *vs.* PM_10_-exposed EAE mice during the pathological course. (**d**) Percentage of disease-free mice after EAE immunization. (**e**) Maximum clinical score reached by EAE mice during the pathological course. (**f**) Cumulative score obtained by saline or PM_10_-exposed EAE mice in 21 days of assessment. Each dot in (**e,f**) represents an individual mouse. Source data are provided as a Source Data file (supplementary material).

**Figure 2.**
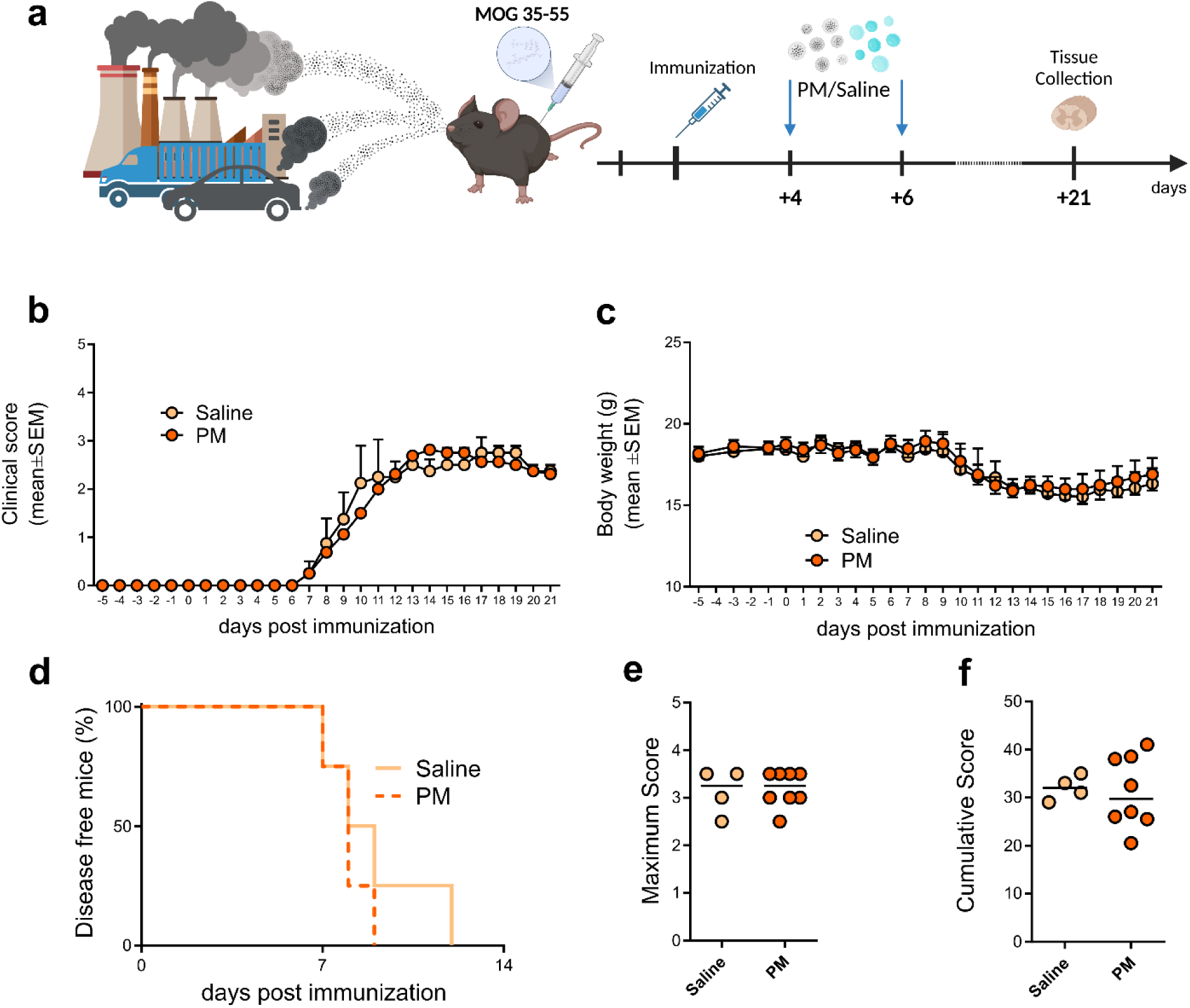
Post-immunization exposure to PM_10_ does not alter EAE pathological course. (**a**) Schematic representation of the experiment design (graphics created with BioRender.com). (**b**) Mean clinical score of saline vs. PM_10_-exposed EAE mice. (**c**) Mean body weight of saline vs. PM_10_-exposed EAE mice during the pathological course. (**d**) Percentage of disease-free mice after EAE immunization. (**e**) Maximum clinical score reached by EAE mice during the pathological course. (**f**) Cumulative score obtained by saline or PM_10_-exposed EAE mice in 21 days of assessment. Each dot in (**e,f**) represents an individual mouse. Source data are provided as a Source Data file (supplementary material).

Consistently, post-immunization exposure to PM_10_ did not affect the neuropathology of the lumbar enlargement of the spinal cord in EAE mice at 21 dpi. Specifically, the extent of demyelination (as assessed by the MOG-negative area - Fig. 3**a**-**c** - and by the quantification of the compact myelin reflectance signal - Fig. 3**d**-**g**), the abundance and hypertrophy of astrocytes (i.e. the GFAP-positive area; Fig. 3**h**-**l**) and microglia (i.e. the IBA1-positive area; Fig. 3**m**-**o**) did not differ in PM_10_- *vs*. saline-exposed EAE mice.

**Figure 3.**
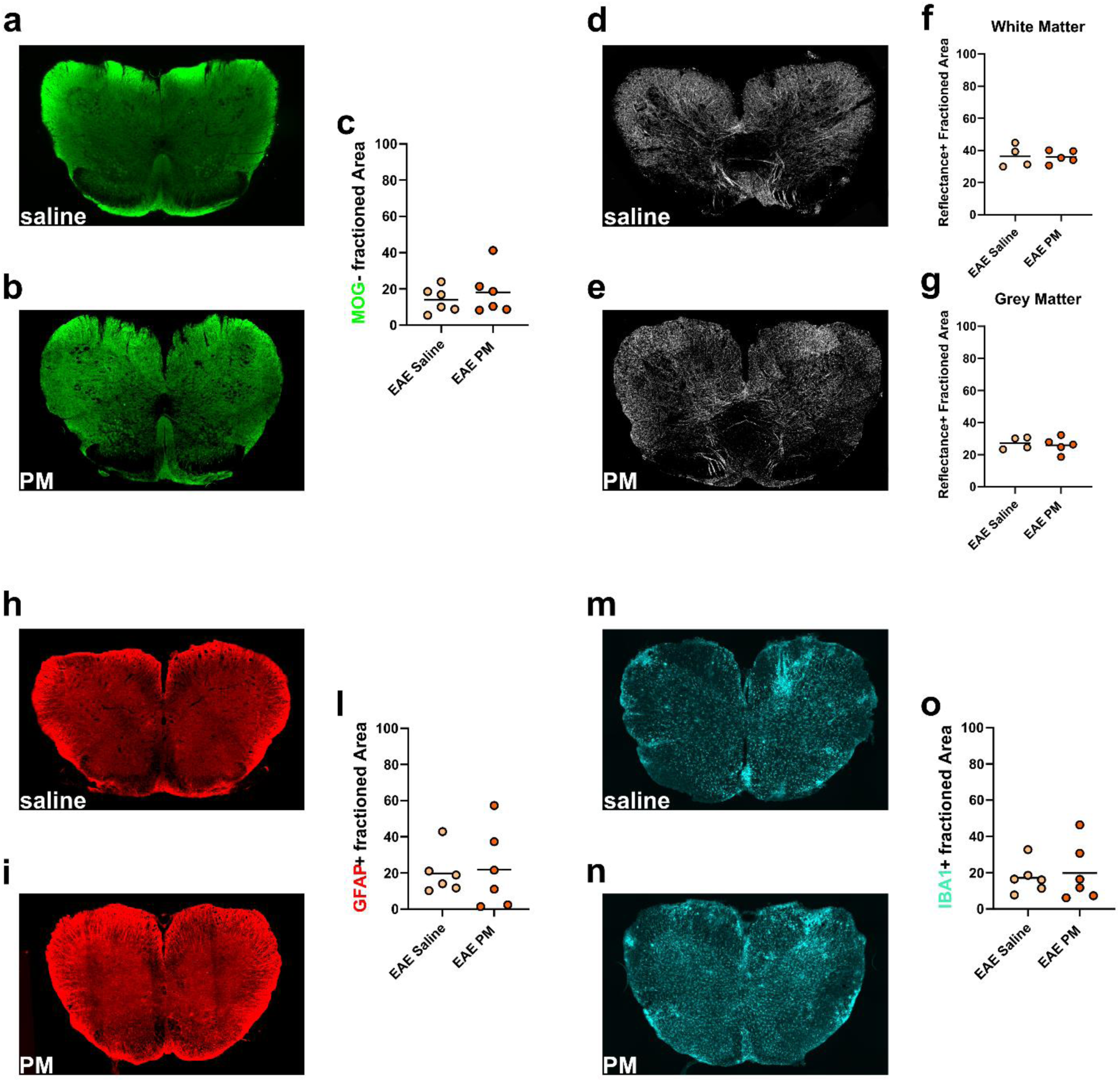
Post-immunization exposure to PM_10_ does not alter EAE neuropathology. (**a,b**) Representative images of anti-MOG immunostaining in saline (**a**) and PM_10_-exposed (**b**) EAE mouse lumbar spinal cord. (**c**) Quantification of the demyelinated (MOG-negative) area. (**d,e**) Representative images of the SCoRe signal (i.e. compact myelin) in saline (**d**) and PM_10_-exposed (**e**) EAE mouse lumbar spinal cord. (**f,g**) Quantification of the SCoRe reflectance signal in the white (**f**) and grey (**g**) matter of the lumbar spinal cord. (**h,i**) Representative images of anti-GFAP immunostaining in saline (**h**) and PM_10_-exposed (**i**) EAE mouse lumbar spinal cord. (**l**) Quantification of GFAP+ area. (**m,n**) Representative images of IBA1 staining in saline (**m**) and PM_10_-exposed (**n**) EAE mouse lumbar spinal cord. (**o**) Quantification of IBA1+ area. Each dot in (**c,f,g,l,o**) represents an individual mouse. Source data are provided as a Source Data file (supplementary material).

### Acute exposure to PM_10_ selectively induces behavioral alterations in presymptomatic EAE mice

In addition to clinical scores and tissue neuropathology – which are the conventional readouts of progressive ascending paralysis in EAE (Montarolo et al., 2015) – we reasoned to assess whether exposure to PM_10_ might influence other variables more predictive of MS co-morbid symptoms, such as fatigue, cognitive impairment and mood disturbances, which are experienced by 60% of persons with MS already in the early phase of the disease (Woo et al., 2024; Benedict et al., 2020; Sparaco et al., 2021). The Open Field (OF) test was performed to evaluate the gross motor function and emotional state (i.e. possible signs of anxiety, which is one of the most common psychiatric co-morbidities in MS patients; Kraeuter et al., 2019; Sparaco et al., 2021) of EAE *vs.* healthy (Ctrl) mice early (i.e. 6h) after PM_10_ exposure (Fig. 4**a-d**). EAE mice exhibited a strong decrease in the total traveled distance (Fig. 4**c**) and mean speed (Supplementary Fig.1**a**), compared to Ctrl mice, in line with an early impairment of motor functions that could not be detected in the conventional clinical score assessment (see above and Fig.2**b-f**). Exposure to PM_10_ did not have a significant effect on these parameters in both EAE and Ctrl mice (Fig. 4**c** and Supplementary Fig.1**a**). Yet, the tracking analysis - which distinguishes the central (i.e. distance traveled in the central part of the arena) *vs*. side (i.e. distance traveled in the “safer” space close to the walls of the arena) paths – detected a significant increase in the preference for the center in EAE mice exposed to PM_10_ compared to Ctrl and saline-exposed EAE mice (Fig. 4**d**), in line with a specific response of EAE mice to PM_10_ manifesting as a risk-taking and disinhibited behavior.

**Figure 4.**
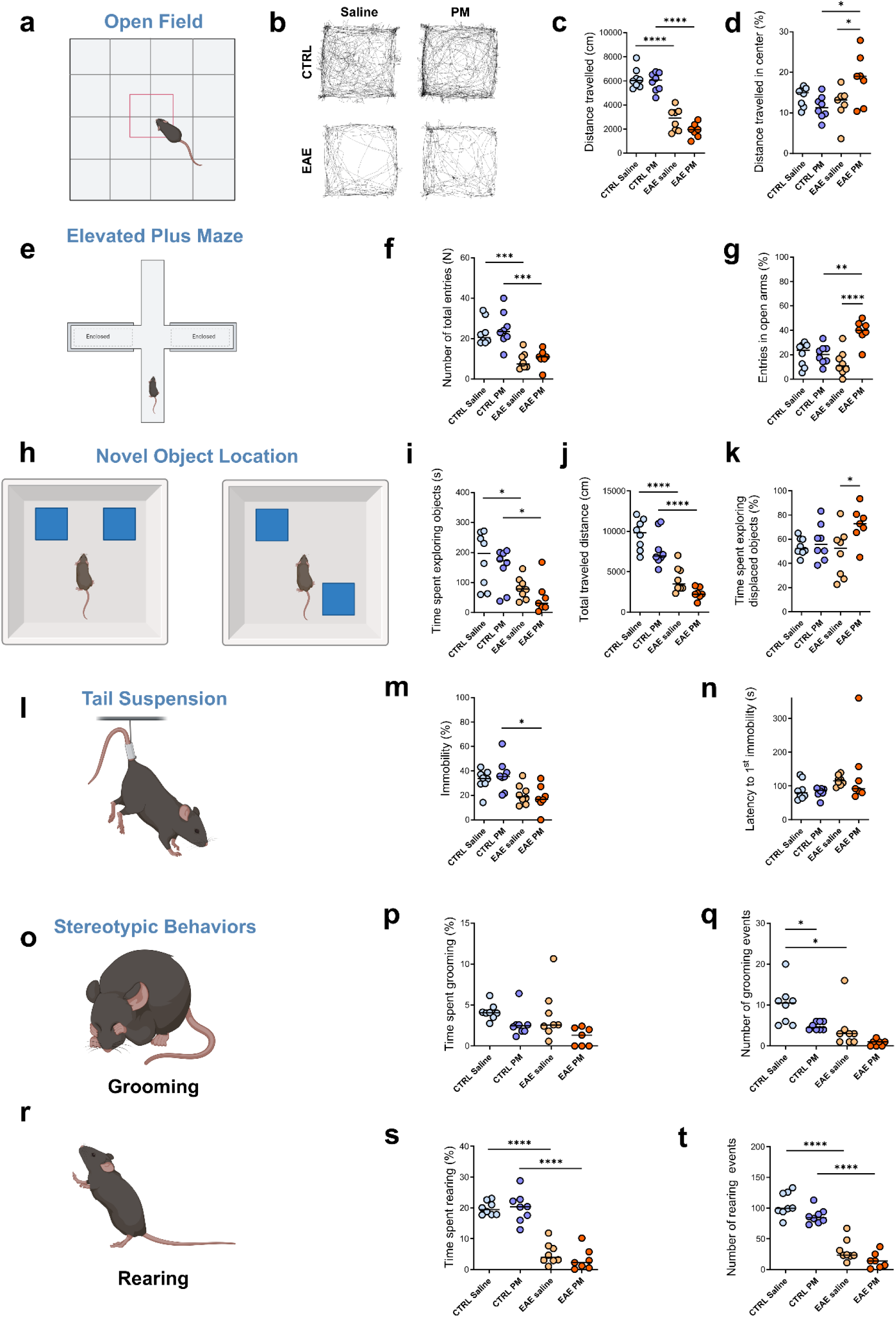
Behavioral alterations elicited in presymptomatic EAE mice after acute PM_10_ exposure. (**a**) Schematic representation of the Open Field (OF) Test. (**b**) Plot tracks of the mouse paths in the OF arena after saline or PM_10_ exposure. (**c**) Total distance traveled by mice during the OF test. (**d**) Percentage of distance traveled in the central part of the OF arena during the test. (**e**) Schematic representation of the Elevated Plus Maze (EPM) test. (**f**) Total number of entries in arms during the EPM test. (**g**) Percentage of entries in open arms during the EPM test. (**h**) Schematic representation of the Novel Object Location (NOL) test. (**i**) Total time spent exploring objects in the NOL test. (**j**) Distance traveled during both trials of the NOL test. (**k**) Percentage of time spent exploring the displaced object in the NOL test. (**l**) Schematic representation of the Tail Suspension test. (**m**) Percentage of time spent in immobility during the TST. (**n**) Latency to first immobility during the TST. (**o**) Schematic representation of the grooming. (**p**) Percentage of total time spent grooming during the OF test. (**q**) Total number of grooming events during the OF test. (**r**) Schematic representation of the rearing. (**s**) Percentage of time spent rearing during the OF test. (**t**) Total number of rearing events during the OF test. Each dot represents an individual mouse. In all instances, differences among groups have been evaluated by means of the Two-ways Anova test followed by Bonferroni’s post-hoc analysis (n, P, and F values in Supplementary Table 1). Statistical differences were indicated with * P *<* 0.05, **P *<* 0.01, ***P *<* 0.001, ****P *<* 0.0001. Source data are provided as a Source Data file. Graphics created with BioRender.com.

To corroborate this interpretation and assess mouse predisposition to risk-taking *vs.* self-protective behaviors, we performed the Elevated Plus Maze (EPM; Fig. 4**e-g**) test, which offers the animals a choice between a risky (open arms) and a safe (i.e. closed arms) area of a four arms apparatus. An increase in mouse activity (duration and/or entries) in open arms reflects an ethologically inconsistent preference for risky options (Walf and Frye, 2007). Consistent with the reduced motility observed in the OF test, both PM_10_- and saline-exposed EAE groups displayed a reduced number of total entries in the arms of the EPM platform (Fig. 4**f**) as well as a reduced traveled distance compared to Ctrl mice (Supplementary Fig.1**b**). Yet, EAE mice exposed to PM_10_ showed an increased number of entries in the open arms of the EPM platform, compared to the other groups (Fig. 4**g**), in line with an increased risk-taking behavior.

We then employed a modified Novel Object Location (NOL) test (in which the inter-trial interval was kept at 10 minutes) to detect shifts in novelty preference (i.e. neophilia, as assessed by increased investigation of novel items; Fig. 4**h-k**), which is important in decision-making and may result in risk-taking behaviors (Swiercz et al., 2025). Approach–avoidance behaviors toward novel stimuli can be affected in a variety of neuropsychiatric disorders, including attention-deficit/hyperactivity disorder, anxiety and depression, whose symptoms are frequently experienced by persons with MS (Sparaco et al., 2021). While both EAE mouse groups moved significantly less than Ctrl mice (as assessed by measuring the time spent exploring objects and the total traveled distance; Fig. 4**i,j**), EAE mice exposed to PM_10_ selectively showed an increased exploration of the displaced object during the second trial of the test (Fig. 4**k**), indicating an increased interest toward novelty.

To explore a possible interaction of PM exposure and EAE in the regulation of emotional behaviors, we performed the Tail Suspension Test (TST), where the duration of immobility (indicative of despair/loss of motivation) *vs.* escape-oriented behaviors are considered as a measure of depressive symptoms (Fig. 4**l-n**; Cryan et al., 2005). While both groups of EAE mice displayed a slight decrease in the percentage of time spent in immobility - maybe due to a higher susceptibility to discomfort (Fig. **4m**) - exposure to PM_10_ did not alter mouse behavior in the TST in both Ctrl and EAE mice (Fig. 4**m**,**n**). Finally, we evaluated mouse spontaneous stereotypic behaviors (i.e. self-grooming and rearing) during free exploration in the OF (Fig. 4**o-t**). In rodents, self-grooming is an innate stereotyped pattern of body cleaning behaviors, whose alterations are recognized as a marker of stress responses and anxiety (Liu et al., 2021). Spontaneous rearing behavior, in which rodents stand on their hindlimbs to explore, can also be quantified in the OF test to provide an additional measure of anxiety (Sturman et al., 2018). Decreased grooming events (Fig. 4**q**) and rearing time/events (Fig. 4**s,t**) were observed in EAE mice, in line with their reduced motility reported in other tests (see above). Yet, exposure to PM_10_ did not trigger any alteration in stereotypic behaviors in both Ctrl and EAE mouse groups (Fig. 4**p,q,s,t**).

Overall, results obtained from mouse phenotypic characterization converge in highlighting a specific vulnerability of EAE mice toward PM_10_-induced behavioral alterations – presenting as increased risk-taking, disinhibited and novelty-seeking behaviors.

**Supplementary Figure 1.**
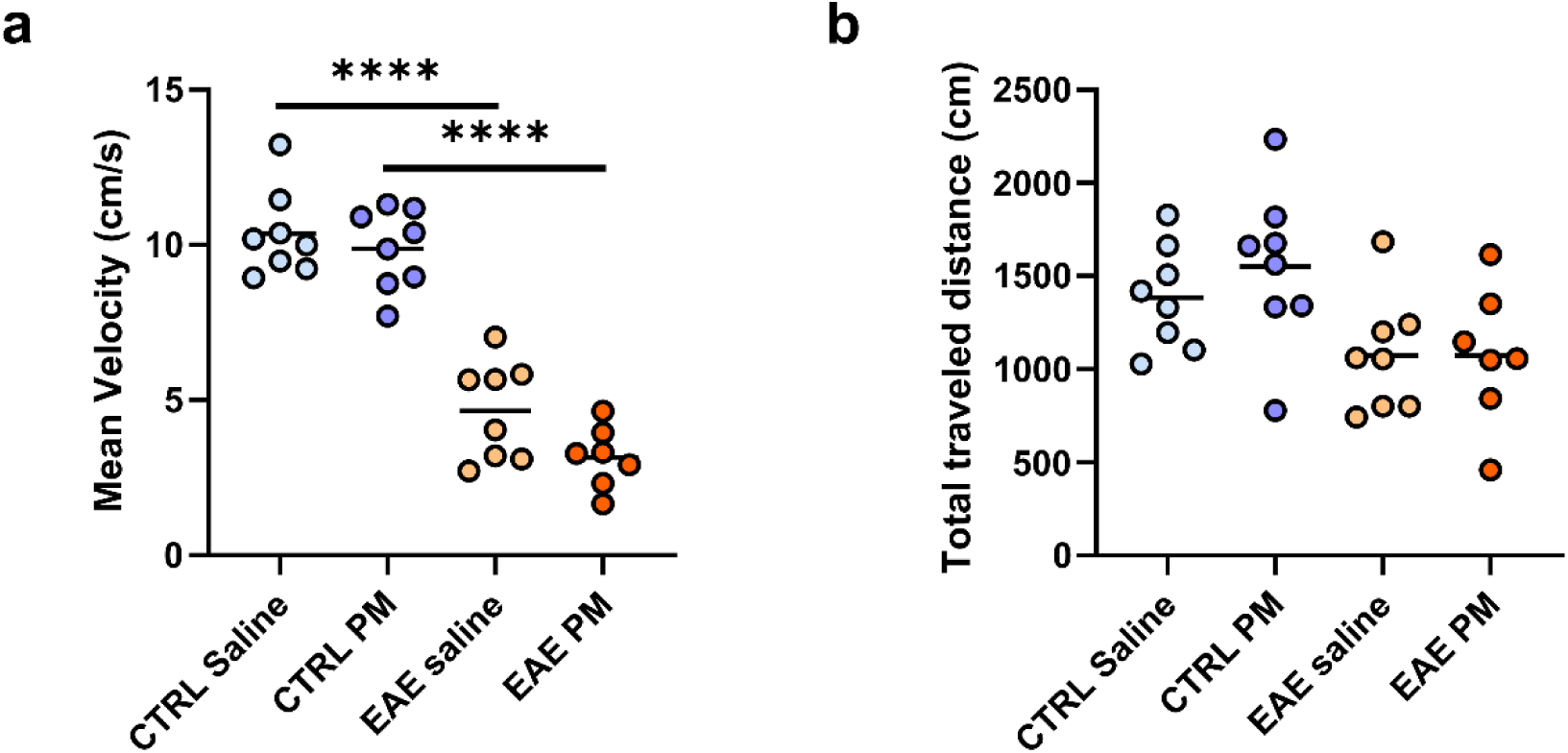
Additional behavioral analyses of Ctrl vs. EAE mice after acute PM_10_ exposure. (**a**) Mean speed of mice during the OF exploration. (**b**) Total distance traveled by mice during the EPM test. Each dot represents an individual mouse. Differences were assessed by Two-way ANOVA (see Suppl. Table 1 for P and F values of each comparison). ****, P<0.001. Source data are provided as a Source Data file (supplementary material).

### Acute exposure to PM_10_ does not alter dopamine availability or dopamine system gene expression in the brain of EAE mice

To assess the biological substrate of the selective response elicited in EAE mice by PM_10_, we first investigated whether PM_10_ induced a different neuroinflammatory response in EAE *vs*. healthy Ctrl mouse brain as early as 6 hours after PM_10_ exposure. Yet, besides an expected increase in INFγ and IL1β in both EAE groups, no significant difference in the gene expression of proinflammatory cytokines (i.e. INFγ, IL1α, IL1β, IL-6, TNF) was found associated with PM_10_ exposure (Fig. 5**a** and Supplementary Fig. 2n-r). In line with this and with the unaltered disease course (see above and Fig. 2**b-f**), exposure to PM_10_ did not accelerate oligodendroglia or myelin loss in presymptomatic EAE mice, as assessed by the quantification of the transcripts coding for the pan-oligodendroglia marker Olig2 and for the myelin basic protein (Mbp; Fig. 5**b** and Supplementary Fig. 2s,t).

**Figure 5.**
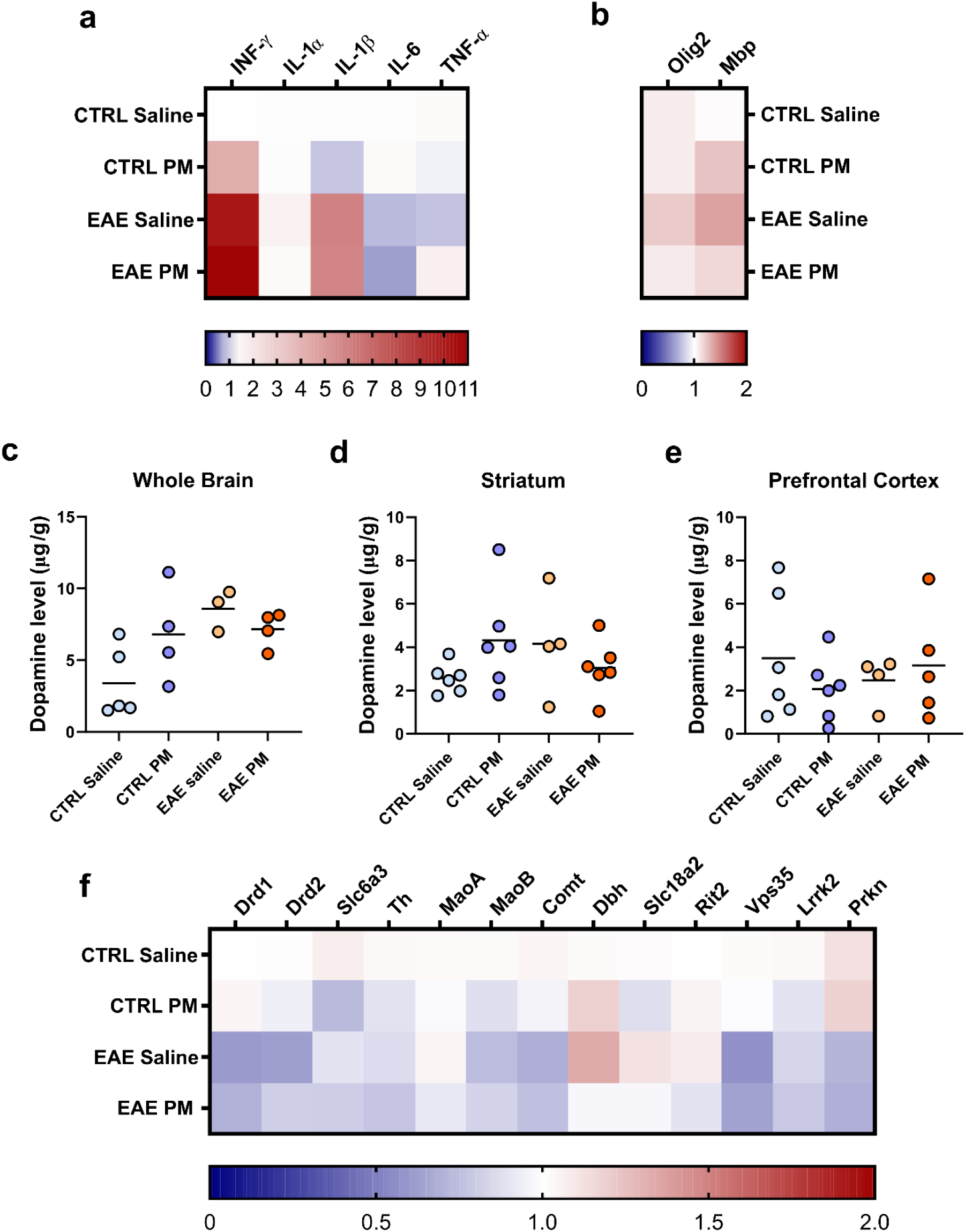
Dopamine availability and expression of dopamine system genes are not altered by exposure to PM_10_. (**a,b**) Heatmap of the qRT-PCR analysis of inflammatory cytokines (**a**) and oligodendroglial genes (**b**) expression. (**c**) Quantification of dopamine by HPLC in the mouse whole brain tissue. (**d,e**) Quantification of dopamine by HPLC-HRMS in dissected mouse striatum (**d**) and prefrontal cortex (**e**). (**f**) Heatmap of the qRT-PCR analysis of dopamine system gene expression. Two-way Anova followed by Bonferroni’s Multiple Comparison Test (n, P, and F values in Supplementary Table 1). Dot plots of the represented data are included in Supplementary Fig 2. Source data are provided as a Source Data file (supplementary material).

Then, we reasoned to investigate possible changes of dopaminergic neurotransmission, which were formerly associated with increased impulsive, risk-taking and novelty-seeking behaviors and alterations in spontaneous locomotion and stereotypic behaviors (Pogorelov et al., 2005; Kalueff et al., 2016; see also Discussion). Brain dopamine levels tended to increase in both EAE groups compared to Ctrl mice as well as in PM_10_-exposed Ctrl mice compared to saline-exposed Ctrl mice, as assessed by HPLC analysis (Fig. 5**c**). Yet, no significant change in dopamine was found associated with PM_10_ exposure in EAE mouse brain (Fig. 5**c**). To assess region-specific alterations in dopamine availability - possibly undetected in our whole brain quantification - dopamine levels have been also investigated in dissected striatum and prefrontal cortex, the two main regions involved in dopaminergic transmission and in the observed behavioral phenotypes (Kim and Lee, 2011; Zeidler et al., 2025). Exposure to PM_10_ did not alter dopamine availability in both areas (Fig. 5**d,e**). We then further looked for possible changes in the expression of genes coding for dopamine receptors, transporters and synthetic/degrading enzymes (Fig. 5**f** and Supplementary Fig. 2), that might be at base of the different behavioral response of EAE mice to PM_10_. Drd1, Drd2 (coding for the dopamine receptors 1 and 2) and Vps35 (coding for the vacuolar protein sorting-35, involved the dopamine transporter endosomal recycling) transcripts appeared decreased in EAE mice, whereas no significant change in gene expression was detected as associated with PM_10_ exposure in both Ctrl and EAE mice (Fig. 5**f** and Supplementary Fig. 2a-m).

**Supplementary Figure 2.**
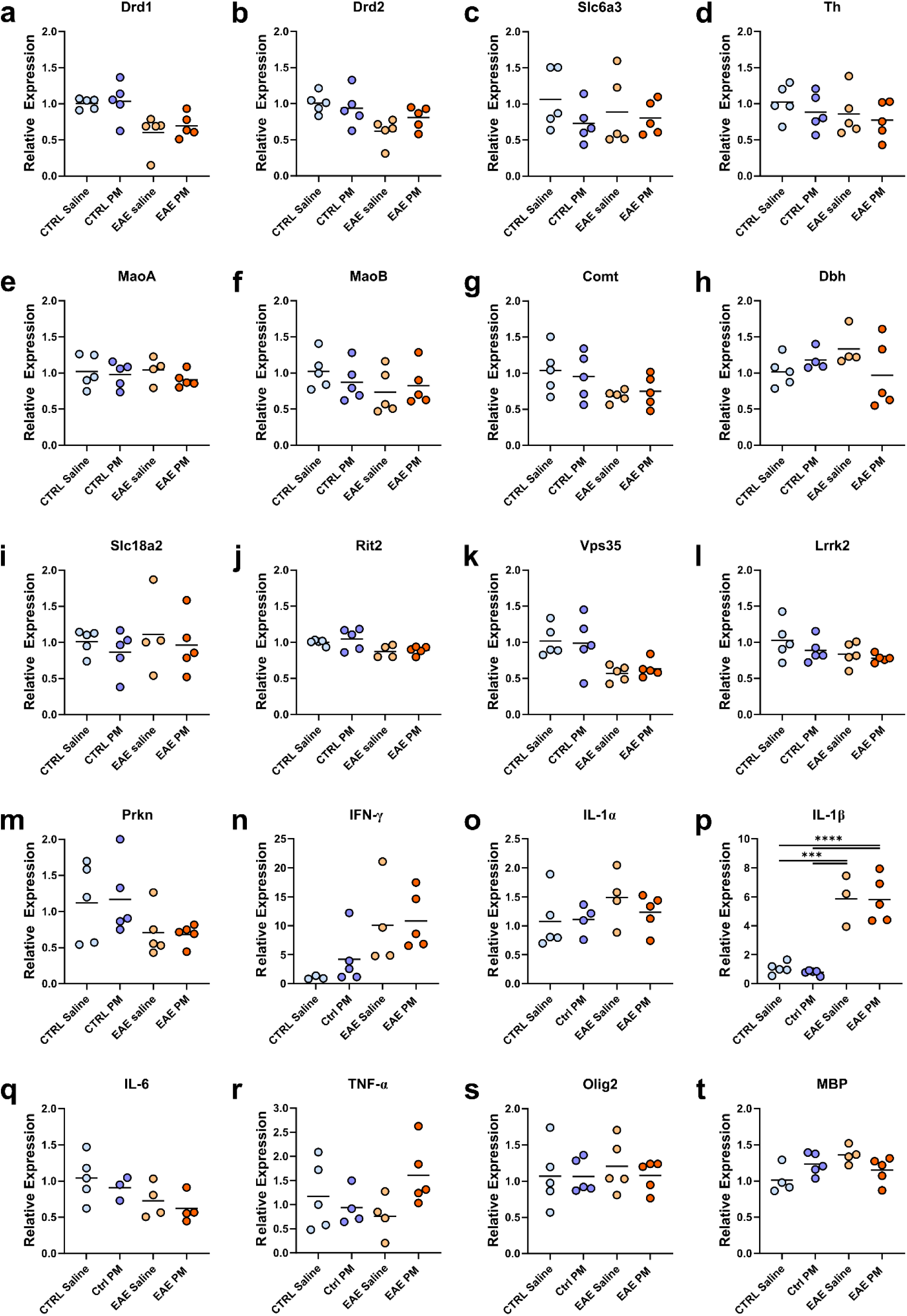
qRT-PCR expression analysis in mouse forebrain. Dot plots referring to Figure 5**a,b,f**. Each dot represents an individual mouse. Differences were assessed by Two-way ANOVA (see Suppl. Table 1 for P and F values of each comparison). ***, P<0.001, ****P *<* 0.0001. Source data are provided as a Source Data file.

## Discussion

Epidemiological studies have shown that transient increases of airborne PM (i.e. PM peaks) are associated with higher rates of hospitalization for MS onset or relapses, and exacerbation of neuroinflammation in MS patients (Angelici et al., 2016; Roux et al., 2017, Heydarpour et al., 2014; Oikonen et al., 2003; Gregory et al., 2008; Noorimotlagh et al., 2021). This prompted the hypothesis that even short-term exposures to high concentrations of PM may contribute to MS emergence in predisposed subjects or trigger relapses in people with MS. To tackle this issue, we combined an immune priming against myelin antigens (i.e. the induction of chronic EAE, the most common animal model of MS; Procaccini et al., 2015) in the mouse with the acute exposure to PM_10,_ either before the immunization or during the pre-symptomatic phase of the EAE course. Although acute PM_10_ exposures did not significantly influence the onset or the severity of EAE in terms of mouse motor impairment (i.e. clinical score), neuropathology and neuroinflammatory/demyelination markers, presymptomatic EAE mice showed behavioral alterations early after being exposed to PM_10_. Specifically, increased disinhibited, risk-taking and novelty-seeking behaviors were triggered 6 hours after exposure to PM_10_ in EAE – but not healthy mice - in line with a specific vulnerability of immunologically primed mice toward the effects of PM well before the emergence of overt functional impairment.

Besides motor, sensory and autonomic disturbances, psychiatric symptoms and cognitive impairment are common in people with MS (Benedict et al., 2020; Sparaco et al., 2021). Psychiatric comorbidity – in most cases in the form of anxiety or depression - is associated with MS, likely due to shared pathophysiological processes, e.g. neuroinflammation (McKay et al., 2018). Additionally, about two-third of MS patients experience cognitive symptoms, including deficits in attention, reduced speed of information processing and executive functioning, and – of relevance for the present findings - cognitive impulsivity and alteration of decision-making abilities (Chiaravalloti and De Luca, 2008; Paul et al., 1998; Toro et al., 2018; Neuhaus et al., 2018; Tabibian et al., 2023). There is increasing evidence and consensus for the importance of the psychiatric and cognitive comorbidities in MS. Presence of these comorbid conditions reduces the quality of personal, social, and professional life and is associated with a significantly higher disability over 10 years of follow-up in persons with MS (McKay et al., 2018). Impulsivity and alteration of decision-making abilities may contribute to poor adherence to therapy, maladaptive coping strategies and engagement in risky behaviors (e.g. tobacco smoking, which is a risk factor for MS disability worsening; McKay et al., 2018). Yet, recognizing and treating cognitive and psychiatric comorbidities in MS may be challenging, as the course of these conditions can be poorly predicted, being detected even in the early stages of MS and influenced by environmental factors. Of note, tobacco smoking - one the major source of indoor PM pollution (Ni et al., 2020) - was recognized as one of the most influential risk factors for worse cognition in early MS (McNicholas et al., 2017). Thus, it would be interesting to assess whether, similar to what was observed in presymptomatic EAE mice, exposure to PM might be associated with increased impulsivity, emergence/worsening of attention deficits or behavior/mood alterations in people with MS, thereby unveiling an additional aspect of vulnerability to air pollution in the cohort of MS patients.

Notably, while most people can homeostatically cope with the negative effects of PM exposure, more vulnerable cohorts exist that show an increased sensitivity toward the consequences of PM exposure. Such susceptibility relies on genetic or acquired factors that make the biological responses triggered by PM different compared to those activated in healthy subjects (Hooper and Kaufman, 2018). The specific nature, timing and selective occurrence of PM-induced behavioral alterations in EAE - but not in healthy - mice, prompted the hypothesis that their biological basis might reside in the interaction of PM-elicited events with an already dysregulated neurotransmitter setting - perhaps consequent to immune priming - in EAE mouse CNS (Akyuz et al., 2023). Among neurotransmitters, we opted for analyzing the possible involvement of the dopamine system for a number of reasons. First, in Dat1 hemizygous mice, reduced expression of the dopamine transporter DAT - the primary mechanism for dopamine clearance after release at synapses - resulted in risk-taking and novelty-seeking behaviors reminiscent of those observed in PM_10_-exposed EAE mice (Pogorelov et al., 2005). Dopamine is also a crucial modulator in the regulation of spontaneous locomotion and stereotypic behaviors, such as grooming (Kalueff et al., 2016). As relevant for MS, dopamine imbalance has been proposed as a substrate of the cognitive fatigue in MS patients (Dobryakova et al., 2015), whereas alterations of dopamine neurotransmission and an imbalance between DR1 *vs.* DR2 receptor signaling were reported as associated with low-grade inflammation in EAE mice (Gentile et al., 2014). Finally, former studies showed that exposure to ambient PM decreased dopamine uptake in the rat striatum (Andrade-Oliva et al., 2023), whereas exposure to PM extracted from tobacco combustion resulted in a rapid decrease of Dat mRNA in the ventral tegmental area and altered DAT function in the dorsal striatum of rats (Danielson et al., 2014). In line with this, in our recent study, dopaminergic neurotransmission was one of the most enriched pathways targeted by EV-packaged miRNAs dysregulated following an acute exposure to PM_10_ in both healthy and EAE mice (Bonato et al., 2025).

Yet, although dopamine tended to increase in the brain of EAE compared to Ctrl mice and in PM_10_- exposed compared to saline-exposed Ctrl mice, no significant change in dopamine was detected in association with PM_10_-exposure in the EAE mouse whole brain tissue. In contrast to our expectations, dopamine levels were unchanged also in dissected striatum and prefrontal cortex, two interconnected areas receiving abundant dopaminergic projections from brain stem nuclei (i.e. substantia nigra and ventral tegmental area) and critically involved in impulse control and threat avoidance in humans and rodents (Kim and Lee, 2011; Zeidler et al., 2025; Fecteau et al., 2007; Liu et al., 2019). Moreover, besides a basal reduction of Drd1, Drd2 and Vps35 mRNAs in EAE vs. Ctrl mice, no significant change in the expression of transcripts coding for dopamine receptors, transporters or biosynthetic/degrading enzymes was detected in either EAE *vs.* Ctrl mice or in PM_10_- exposed *vs.* saline-exposed EAE mice.

These findings suggest the involvement of alternative mechanistic substrates. The most obvious candidate is serotoninergic neurotransmission. Pharmacological studies and genetic manipulations show that serotonin (5-hydroxytryptamine or 5-HT) plays a critical role in risk assessment and impulsivity in humans and animal models. Low serotonin levels have been linked to increased impulsivity and risk-taking behaviors, whereas higher serotonin levels/activity have been associated with increased activation of brain regions involved in risk assessment, e.g. the prefrontal cortex and the amygdala, and to increased risk aversion (Blanchard and Meyza, 2019; Colwell et al., 2024). For instance, Tph2-deficient mice, carrying a germinal deletion of the rate-limiting serotonin-producing enzyme Tryptophan hydroxylase 2 (Tph2), show an impulsive phenotype (Angoa-Pérez et al., 2012; Mosienko et al., 2012, 2015). In line with this, dysregulation of serotonin 1B (5-HT1BR) or 2A (5-HT2AR) receptors can lead to increased impulsivity and preference for risky decisions (Nautiyal et al., 2015; Brunner and Hen, 1997; Macoveanu et al. 2013). Of note, decreased levels of serotonin have been reported in MS patients, likely due to an alteration of tryptophan metabolism induced by inflammatory cytokines (San Hernandez et al., 2020). A second candidate mechanism possibly underlying behavioral alterations observed in EAE mice following exposure to PM_10_ is the neuromodulatory endocannabinoid system (Jacob et al., 2009). In line with this idea, higher levels of anandamide (AEA) and increased expression of CB_1_ and CB_2_ receptors have been reported in the nervous tissue of EAE mice and MS patients (Centonze et al., 2007; Chiurchiù et al., 2018). Whether exposure to PM_10_ may elicit a dysregulation of these systems or operates on an already dysregulated neurotransmitter setting in EAE mice deserves further investigation.

In conclusion, our study unveils a specific vulnerability of EAE mice toward PM_10_-induced behavioral alterations even before they manifest overt neuropathological signs, in line with the idea that the immunological background of the subjects influences the outcome of PM_10_ exposure and pointing to immunologically primed individuals - such as people with MS - as a vulnerable population cohort toward the effects of air pollution.

## Supporting information

Souce Data File

## Author contributions

MB, FM: Data curation, Formal analysis, Investigation, Methodology, Visualization, Writing – original draft, Writing –review & editing.

RP, MC, GA, VC, GN, NDC, CP, VS, FL, SDF: Data curation, Formal analysis, Investigation, Methodology, Writing –review & editing.

MG, AB, AB: Conceptualization, Writing –review & editing.

EB: Conceptualization, Data curation, Formal analysis, Visualization, Writing – original draft, Writing – review & editing, Funding acquisition, Project administration, Supervision.

## Data availability

All data are available in the main text or in the Source data file enclosed as supplementary material.

## Acknowledgements

We wish to thank Dr. Marco Cambiaghi (Department of Neuroscience, Biomedicine and Movement Sciences, University of Verona, Italy) for precious help in NOL test analysis. HPLC-HRMS was performed at the Atlantis Centre for Research on Inflammation and Metabolism in Cancer and in Chronic Degenerative Diseases (https://www.atlantis.unito.it/en). Our work was supported by FISM - Fondazione Italiana Sclerosi Multipla (Italy) (ID:2019/PR-Multi/003) and by Cassa di Risparmio di Torino (CRT) Foundation grant (ID: 2021.0657) to EB. MB was supported by a PON R&I 2014-2020 PhD Fellowship “Dottorati di ricerca su tematiche green e dell’innovazione”, financed by Ministero dell’Istruzione, dell’Università e della Ricerca—MIUR (Italy) in the frame of FSE– REACT EU. This study was also supported by Ministero dell’Istruzione, dell’Università e della Ricerca—MIUR (Italy) project “Dipartimenti di Eccellenza 2018–2022” and “Dipartimenti di Eccellenza 2023–2027” to Dept. of Neuroscience “Rita Levi Montalcini” of the University of Turin.

## List of abbreviations

5-HT: 5-hydroxytryptamine (Serotonin)
AEA: Anandamide
AOBS: Acoustic-optical Beam Splitter
BBB: Blood Brain Barrier
CNS: Central Nervous System
Comt: Catechol-O-Methyltransferase
CSF: Cerebrospinal Fluid
Ctrl: Control DA - Dopamine
DAPI: 4,6-diamidino-2-phenylindole
Dbh: Dopamine Beta Hydroxylase
DDA: Data Dependent Acquisition
DRD1: Dopamine Receptor D1
DRD2: Dopamine Receptor D2
EAE: Experimental Autoimmune Encephalomyelitis
EPM test: Elevated Plus Maze Test
EVs: Extracellular Vesicles
FTMS: Fourier Transformer Mass Spectrometry
GFAP: Glial Fibrillary Acid Protein
HPLC: High Performance liquid Chromatography
HRMS: High Resolution Mass Spectrometry
IBA1: Ionized Calcium-binding Adapter molecule 1 IL-1α - Interleukin 1 alpha
IL-1β: Interleukin 1 Beta IL-6 - Interleukin 6
INF-γ: Interferon Gamma
LOD: Limit of Detectable
LOQ: Limit of Quantification
Lrrk2: Leucine-rich Repeat Kinase 2
MaoA: Monoamine Oxiade A
MaoB: Monoamine Oxidase B
MBP: Myelin Basic Protein
MOG: Myelin Oligodendrocyte Glycoprotein
MRI: Magnetic Resonance Imaging
MS: Multiple Sclerosis
MS/MS: Tandem Mass Spectrometry
NOL test: Novel Object Location test
OF test: Open Field test
Olig 2: Oligodendrocyte Transcription Factor 2
PB: Phosphate Buffer
PBS: Phosphate Buffer Saline
PFC: Prefrontal Cortex
PM: Particulate Matter
Prkn: Parkin RBR E3 Ubiquitin Protein Ligase
qRT-PCR: quantitative Real Time Polymerase Chain Reaction
Rit2: Ras-like without CAAX 2
ROI: Region of Interest
RP column: Reverse Phase column
RT: Reflectance/Transmission
SCoRe: Spectral Confocal Reflectance Microscopy
SE: Standard Error
Slc18a2: Solute Carrier family 18 member 2
Slc6a3: Solute Carrier family 6 member 3
SPE column: Solid Phase Extraction column
Th: Tyrosine Hydroxylase
TNF-α: Tumor Necrosis Factor Alpha
TOF: Time of Flight
Tph2: Tryptophan hydroxylase 2
TST: Tail Suspension Test
VIP-HESI: Vacuum Insulated Probe Heated Electrospray Ionization
Vps35: VPS35 Retromer Complex Component

---

**Supplementary Table 1.**
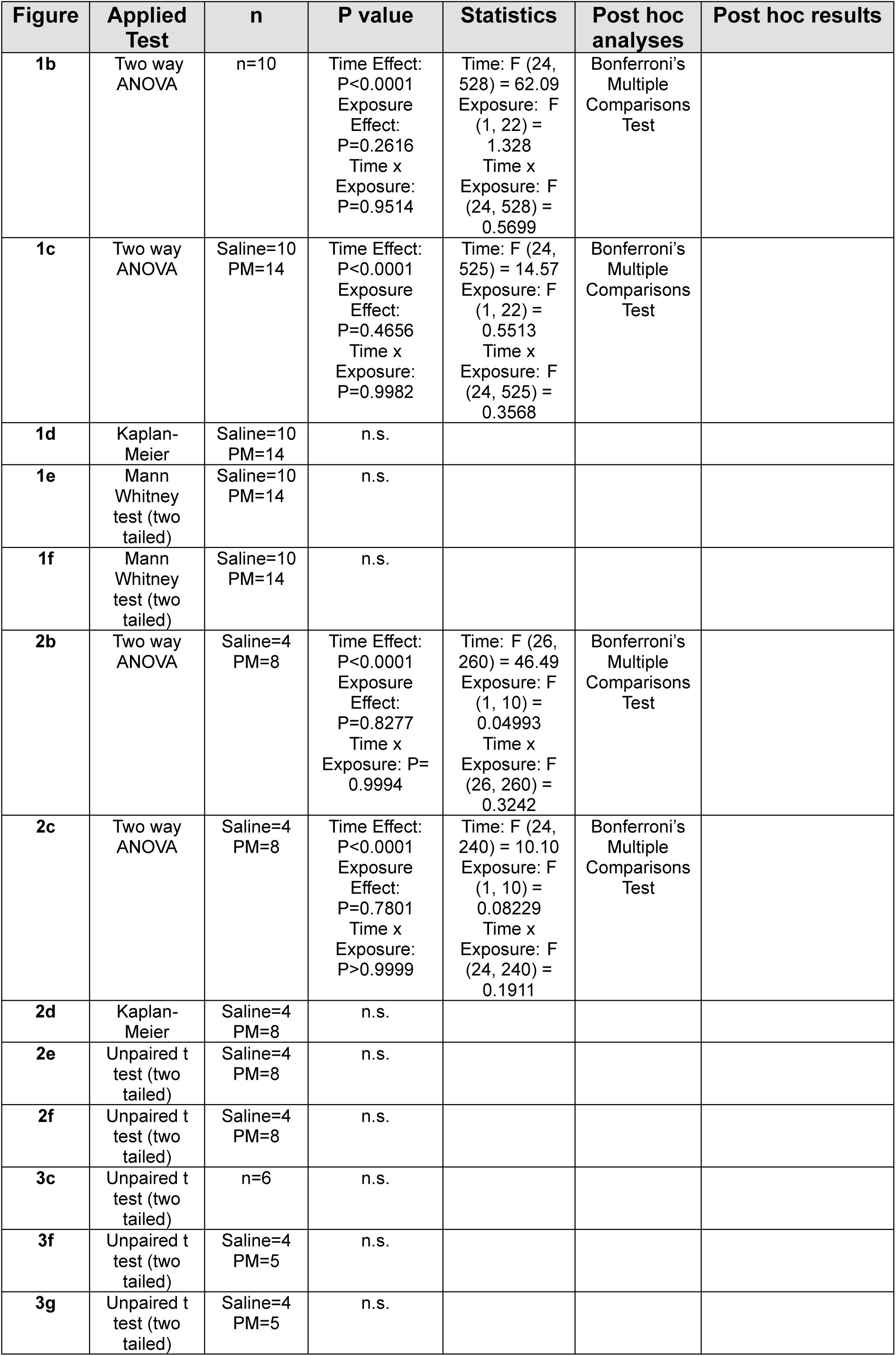
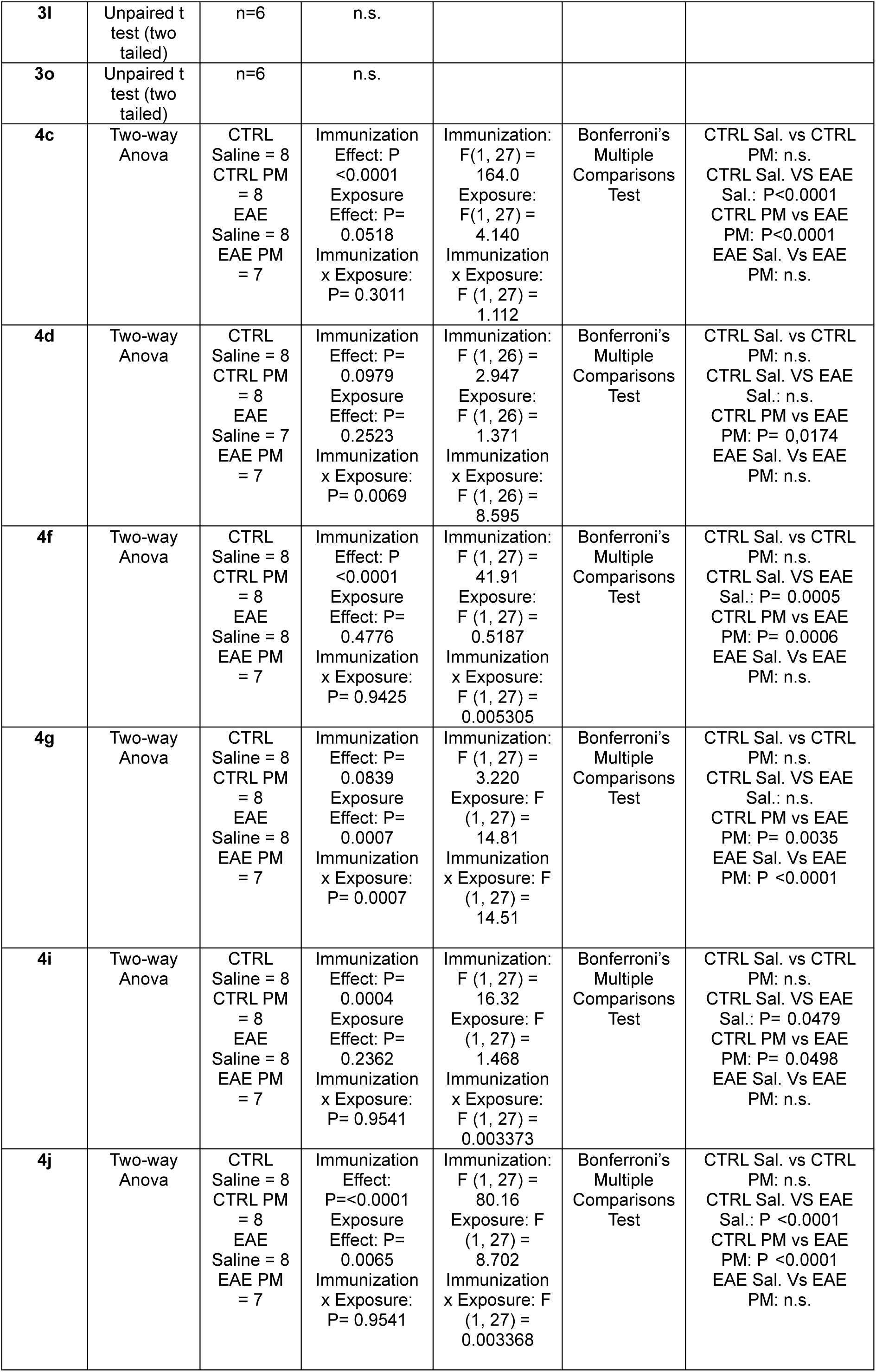
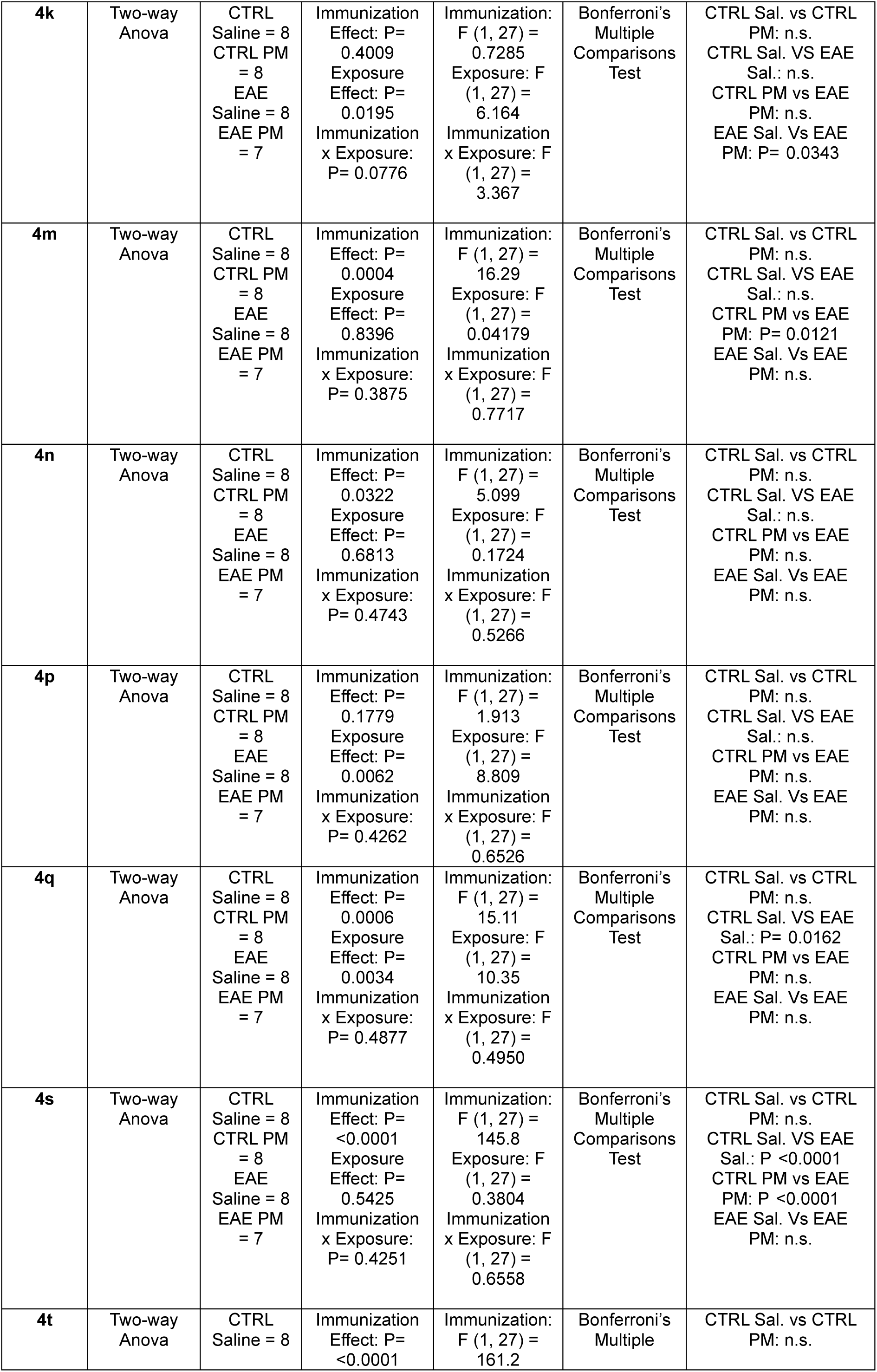
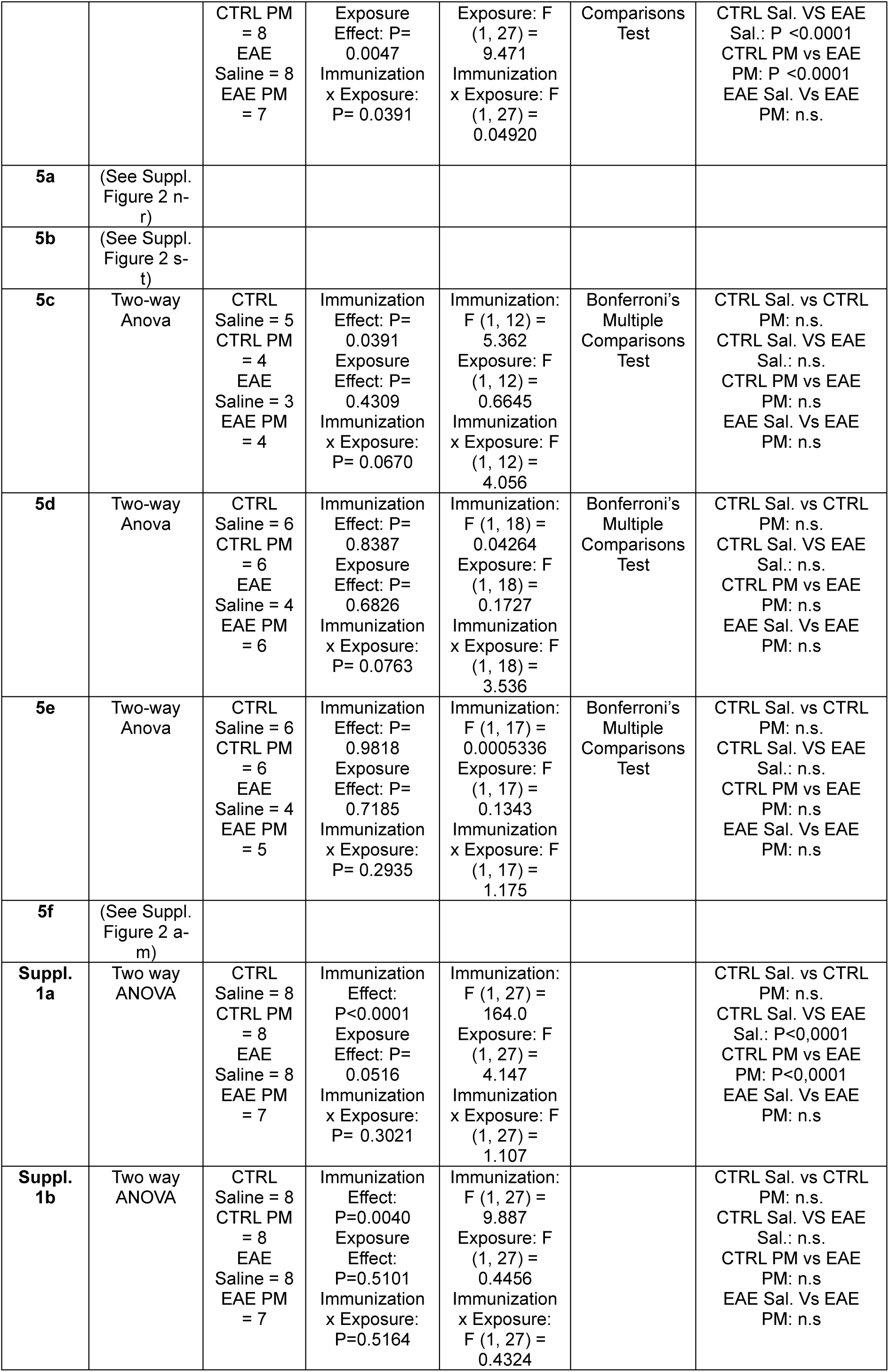
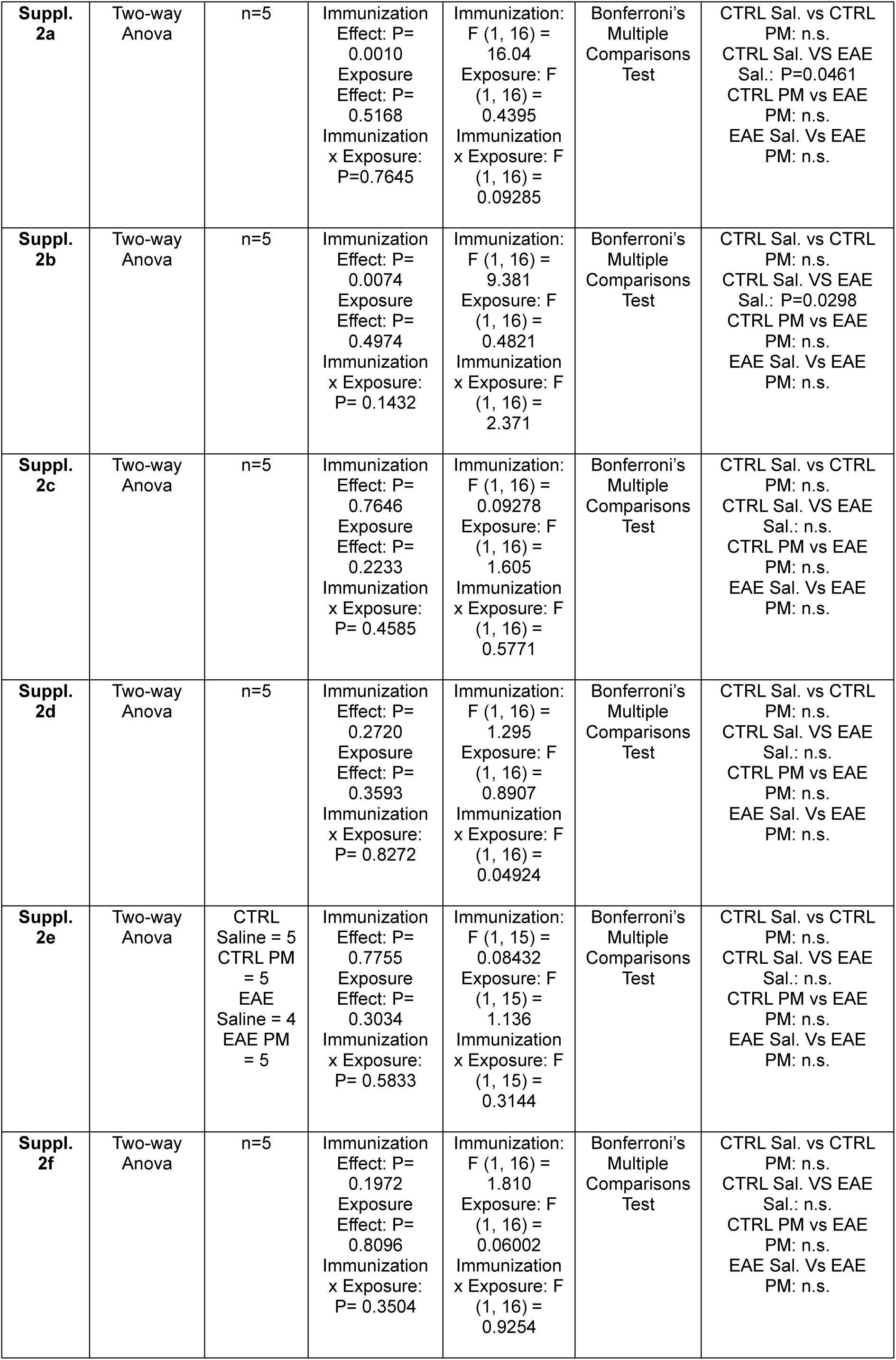
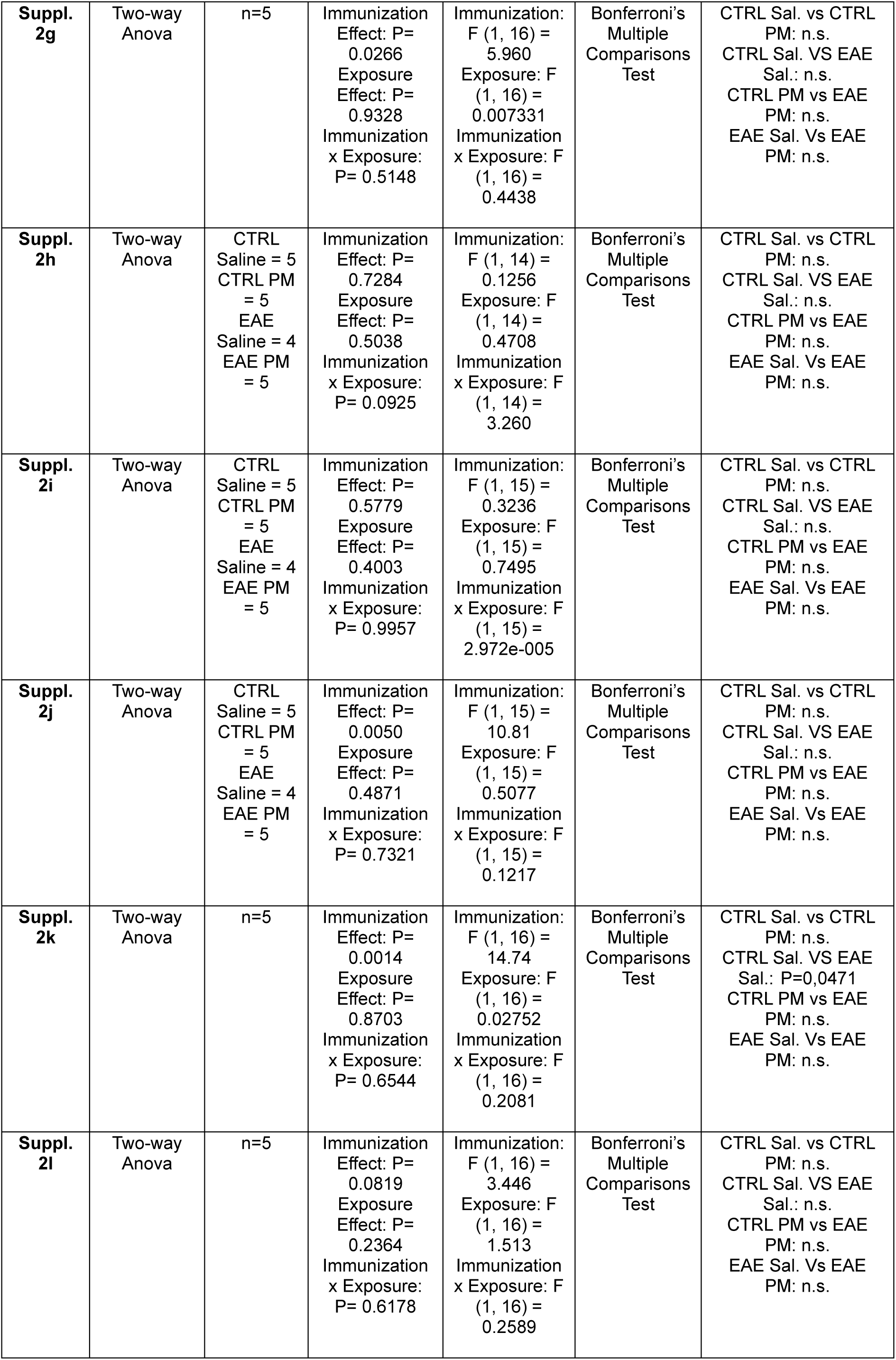
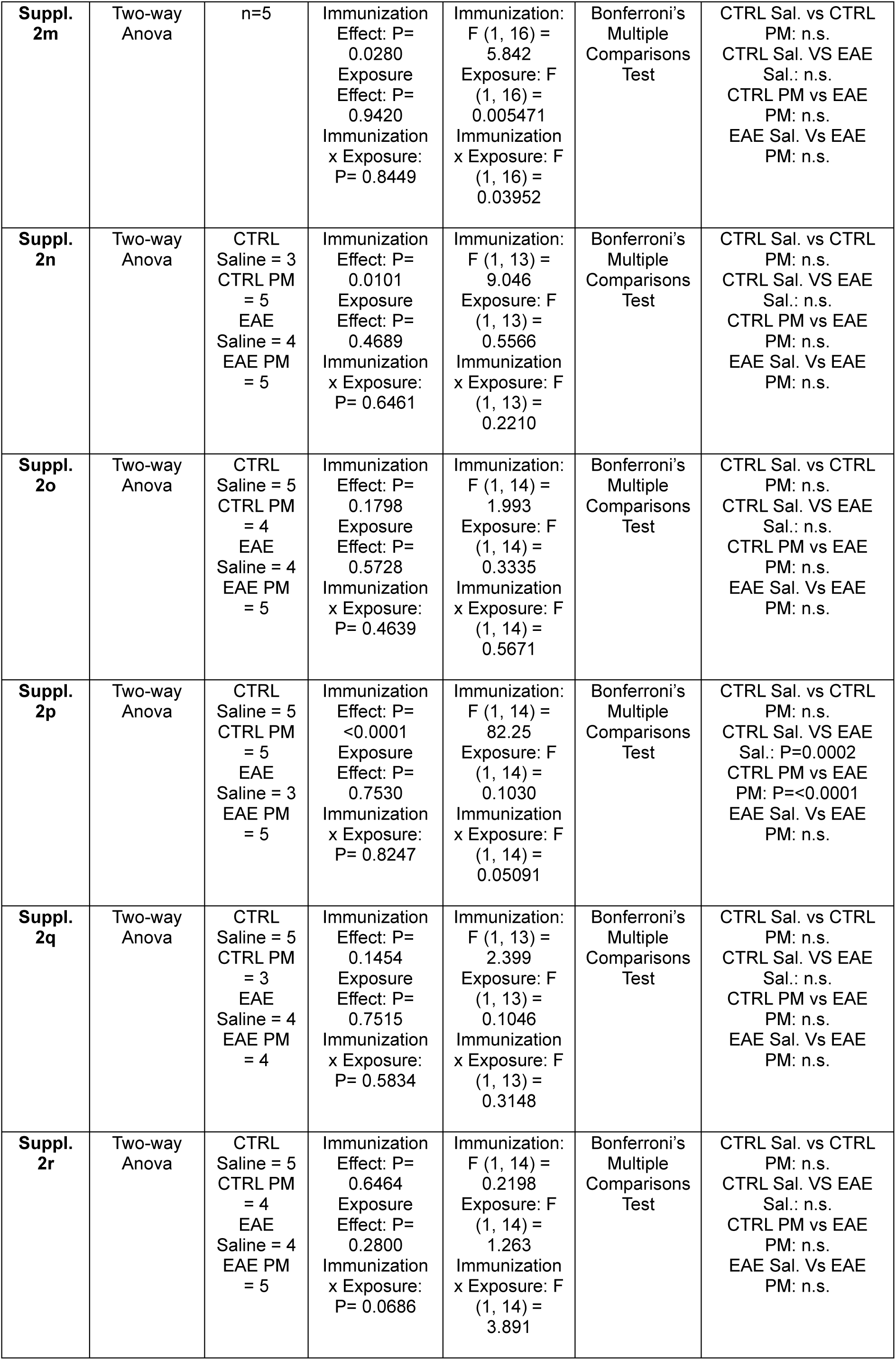
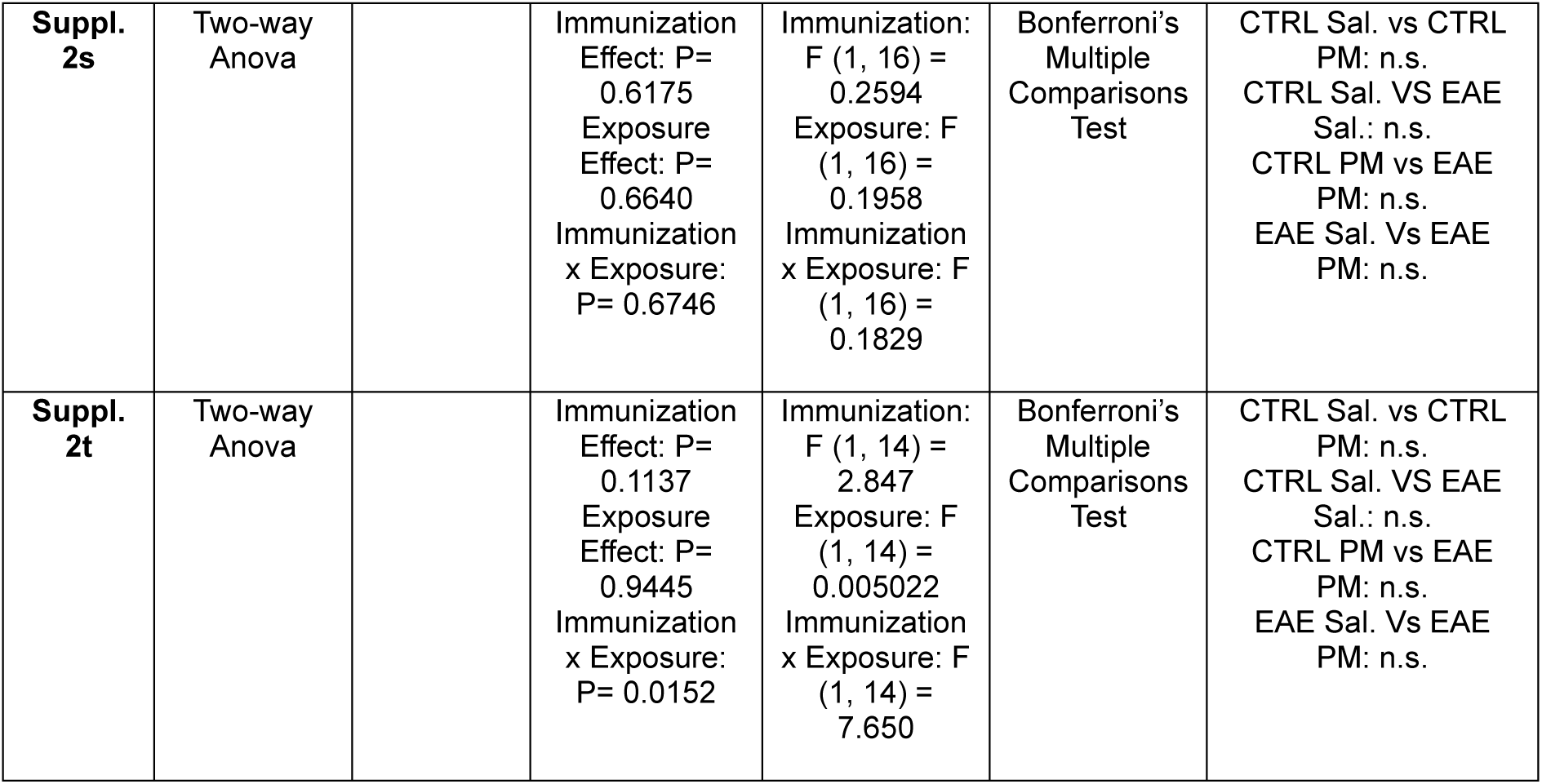
Statistical Analyses.

**Supplementary Table 2.**
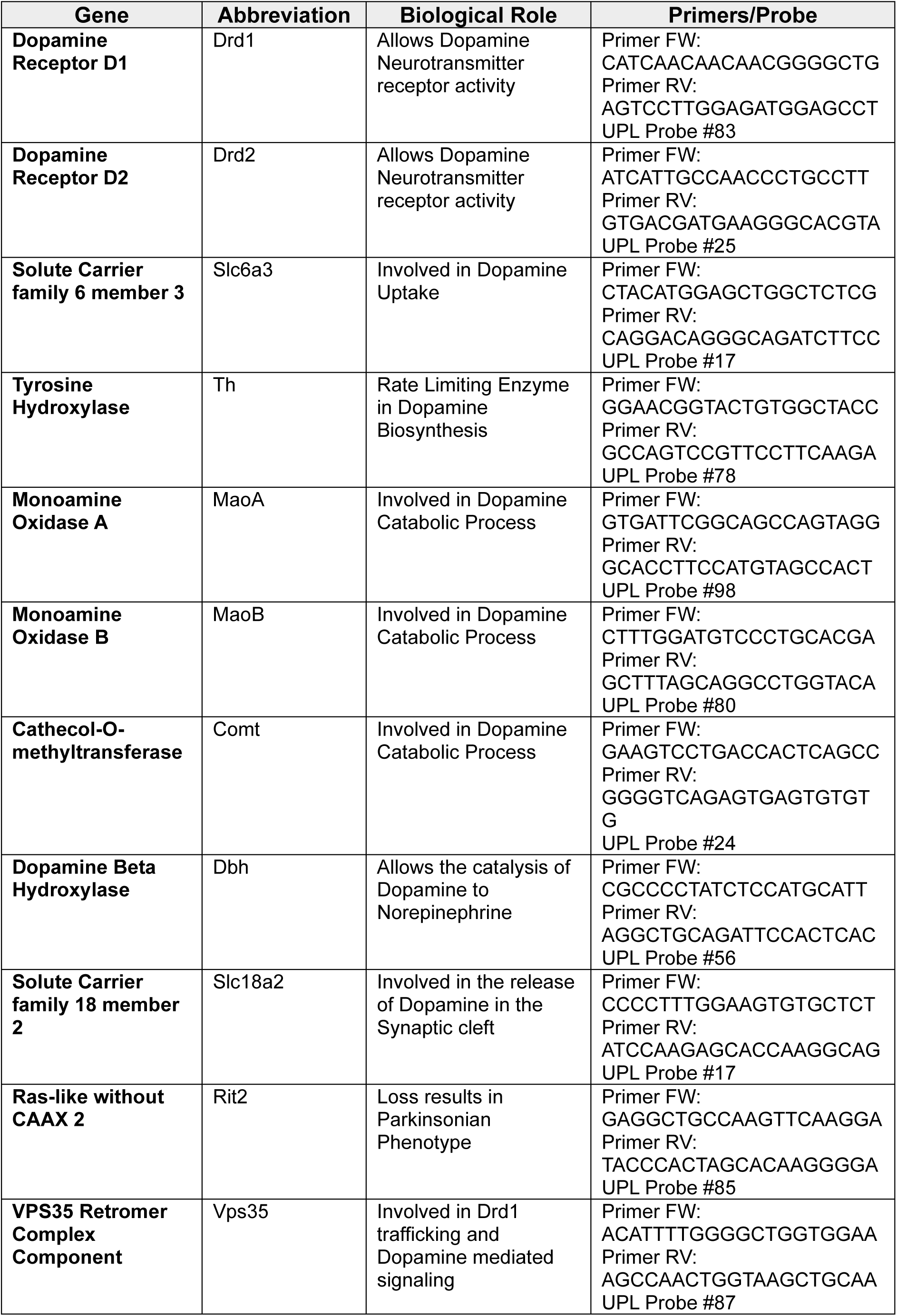
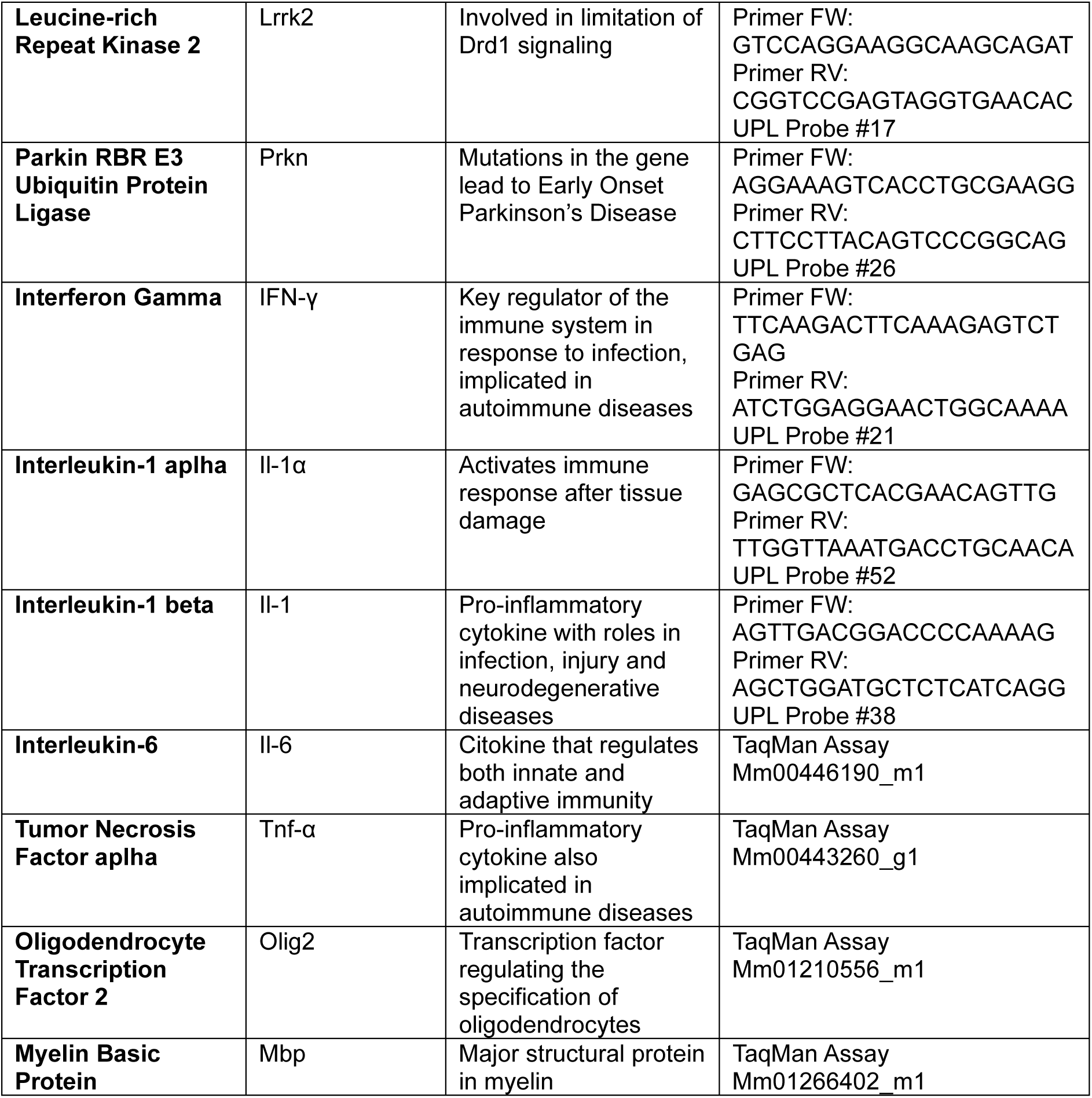
qRT-PCR Assays.

